# Incorporation of single-neuron projectome-based connectivity motifs enhances the cortex-specific performance of artificial neural networks

**DOI:** 10.64898/2026.06.12.732007

**Authors:** Yue Sun, Wangzi Yao, Jian Zhang, Wei Song, Xuanle Zhao, Chonghe Hao, Xinyu Chen, Shuxin Zeng, Shuncheng Jia, Yun Yang, Xu Chen, Xiong Xiao, Mu-ming Poo, Yangang Sun, Bo Xu, Tielin Zhang

## Abstract

The organizational principles of natural neural networks could inspire the new architecture design of artificial neural networks (ANNs). Analysis of single-neuron connectomes of mouse brains revealed distinct profiles of three-node connectivity motifs in various cortical areas and hippocampal formation. A connectome-informed neural network algorithm (“CINA”) was developed to incorporate natural connectivity motifs into ANN algorithms represented by recurrent neural network (RNN) and transformer-based large language model (LLM). We found that incorporation of the average profile of cortical motifs improved the RNN’s performance in noise-resistant categorization and motor learning benchmark tasks, as compared with RNNs with random connectivity. Notably, incorporating cortex-specific motifs further elevated the RNN’s performance in tasks related to the cortical function, and this effect was enhanced by artificially increasing the bias in the motif profile. Similar experimental results were verified on an LLM using Motif-Transformer for natural language question answering and brain-signal decoding tasks. Graph-theoretic analyses showed that incorporating natural motifs drove the emergence of modular and small-world properties in ANNs. Together, we demonstrated not only connectome-inspired optimization of ANN architecture but also functional significance of specific motif profiles in various cortices.

## INTRODUCTION

Cognitive functions of the brain emerge from the complex connectivity among neurons. Recent single-neuron connectomic analyses have begun to provide information on both long-range projections and local connections of neurons in various brain regions^1,2^. Studies of artificial neural networks (ANNs) have also shown that the efficiency of ANNs could be improved by mimicking the architecture of brain circuits. For example, local receptive fields of the visual cortex have inspired the Gabor filter design of signal processing in ANNs^3^, and long-short-term memory networks^4^ were developed to mimic gate-based recurrent connections of memory-attention pathways. In principle, incorporating biological circuit architectures into ANNs may enhance the capability of ANNs towards brain-like intelligence and offer new insights into the structure-function relationship of neural circuits in the brain. However, most artificial network designs have not considered the incorporation of specialized connectivity patterns of the brain, including, but not limited to, recurrent neural networks (RNNs) and the Transformer that serves as the key module in large language models (LLMs).

Selective connectivity among a small group of neurons, known as “motifs”^5^, have been identified and characterized in various brain regions^6^. For example, feedforward and feedback inhibition motifs are employed to regulate the dynamic range of input signals, which play an important role in maintaining excitation–inhibition (E–I) balance across the network, thereby improving the speed and signal-to-noise ratio of information processing^7,8^. Furthermore, the frequencies of certain connectivity motifs are significantly higher than those based on random network connectivity, suggesting specific functional roles of connectivity motifs in the brain^9^. In this study, we inquired whether incorporating distinct distributions of various connectivity motifs derived from different mouse brain regions could improve the performance of RNN and Transformer in machine learning tasks.

We first analyzed recently collected connectome data on the mouse brain (https://www.digital-brain.cn/) for the distribution of 3-node motifs in 50 brain regions. Previous methods for determining the frequency distribution of motifs heavily depended on time-consuming sampling techniques and limited primarily to statistical analyses^10–12^. In this study, we developed a connectome-informed neural network algorithm (termed “CINA”). This method uses matrix multiplication to parallelize the calculation of frequency distributions for all 13 types of three-node motifs. It transforms pure statistical analyses into computable equations that are then integrated into ANN algorithms via a specially designed motif loss function. This newly added loss function serves as a structural constraint that complements the traditional error loss. While the traditional error loss drives the network toward computational accuracy, this additional loss function enforces biological plausibility in terms of network structure during the learning process.

We then examined the benefits of CINA-endowed motif distributions in both RNN and Transformer algorithms for a variety of machine learning tasks, including four types of noise-resistant categorization, four types of reinforcement-based motor learning, and six standard language benchmarks. We found that the performance of RNNs was enhanced by incorporating an average motif distribution found in the frontal pole cortex (FRP) and primary motor cortex (MOp), as compared to that of RNNs with random connectivity. Moreover, incorporating FRP- and MOp-specific motif distributions led to further enhancement of the tasks related to these two cortices. Notably, such selective enhancement by cortex-specific motifs could be further elevated by artificially increasing the cortex-specific motif distribution bias. Similar conclusion could be given for motif-informed Transformer in natural language question answer and brain signal decoding tasks. Finally, we applied graph-theoretic analysis to ANNs endowed with cortical motif distributions and found that they exhibit modular and small-world network properties inherent in neural networks of the brain. Taken together, these findings demonstrated that the performance of ANNs could be enhanced by incorporating biological connectome-based motif distributions into their network architecture.

## RESULTS

### Motif distributions in mouse cortices based on single-neuron connectomes

As the first step, we used the single-neuron whole-brain connectome dataset (https://www.digital-brain.cn/, containing 50 mouse brain regions) which was obtained by sparse labeling of cortical neurons, imaging with the fluorescence Micro-Optical Sectioning Tomography (fMOST), single-axon tracing, and registration to the standard mouse brain template^13^ (**Fig. 1a**). The putative pre- and postsynaptic sites were determined to infer neuronal connectivity. As illustrated in **Fig. 1b**, when an axonal arbor of source neuron falls within a specific distance (e.g., 5 μ*m*) of a dendritic arbor of target neuron in 3D, the existence of a putative site for synaptic contacts from source to target neurons was identified with a connection score. The total connection score is calculated by summing the scores of all putative synaptic sites (see Methods). A matrix of connection scores among all neurons within a cortical region was constructed, as illustrated using data from MOp (**Fig. 1c, left**). Based on the matrix and a preset pruning threshold value of connectivity score, the connectivity among all neurons was obtained (**Fig. 1c, right**). For example, the existence of projection was identified when the connection score is above the threshold. We further identified all 3-node connectivity motifs, which comprise 13 possible configurations, as illustrated by examples from MOp dataset (**Fig. 1d**).

**Fig. 1.**
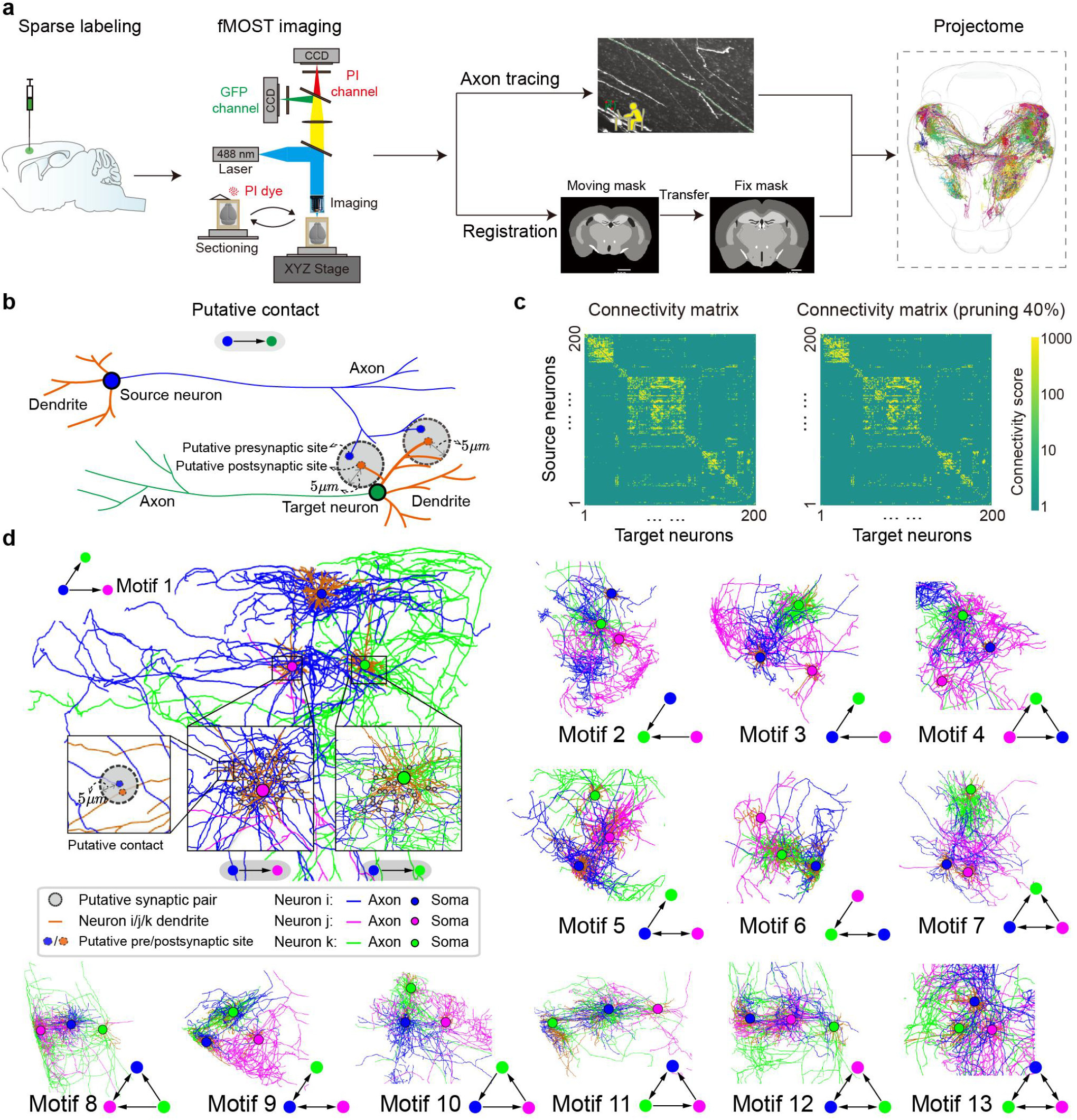
Inference of connectivity motifs from single-neuron mouse projectomes. **a**, Diagram showing workflow for single-neuron whole-brain connectome data collection including sparse labeling, fMOST imaging, axon tracing, and registration. **b**, Schematic diagram depicting the determination of putative pre- and postsynaptic sites used for inferring connectivity. The axon of the source neuron (blue) and the dendrite of the target neuron (orange) are sampled along their arbors (blue and gray points, respectively). When any axonal arbor falls within a threshold (dashed circle) of a dendritic arbor in 3D, a putative connection is registered. **c**, Schematic diagram of the connectivity matrix calculated from **b. d**, Illustration of 13 possible three-node motifs based on single-neuron connectome data from the mouse primary motor cortex (MOp). Single-neuron tracing of axonal/dendritic arbors are projected to the same template, and putative synaptic contacts among three neurons were determined by measuring Euclidean distances between axonal and dendritic arbors of two neurons (see Methods). Example connectivity patterns for each of 13 three-neuron motifs were shown. Colored circles, examples of putative synaptic contacts (for neuron *i* → *j* and *i* → *k*). Arrows, direction of functional projections (connectivity score above a defined threshold).

Based on mouse connectome dataset, we obtained the frequency distribution of 13 three-node motifs in 50 brain regions. The normalized *Z* score^5,9^ was used here to quantify the frequency of a specific motif that differed from that found in a randomly connected network (+, above; −, below; see Methods). Notably, we found that the frequency of fully recurrent three-node motif (type 13) was consistently elevated across all brain regions and frequency distributions were distinctly different among brain regions related to cognitive, motor, and sensory functions (**Fig. 2a**). Further analysis of the similarity in the pattern of motif distributions among 50 brain regions (see Methods) revealed four major clusters corresponding roughly to PFC, hippocampal formation, motor-sensory cortices, and audiovisual cortices (**Fig. 2b)**.

**Fig. 2.**
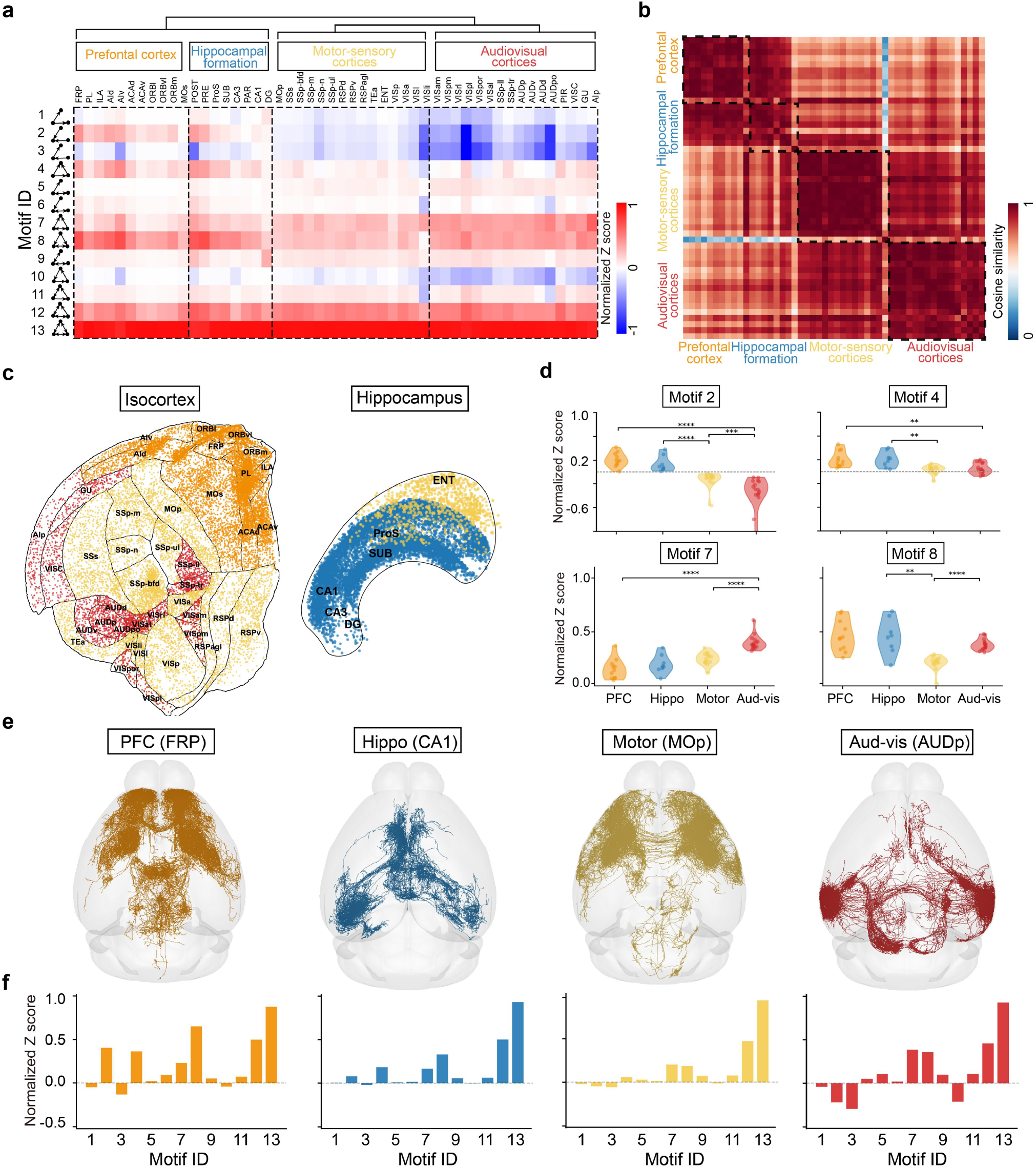
Connectivity motif signatures across 50 mouse brain regions. **a**, Heatmap showing the frequency distribution of 13 three-node motifs relative to random networks for 50 brain regions (names are listed in **Table S1**). Color intensity codes the normalized *Z* score for the deviation from random distribution (red and blue, higher and lower than random, respectively). The frequency of the fully recurrent three-node connectivity motif (type 13) was consistently higher than that of random networks across all brain regions. Frequency distribution patterns were distinctly different among brain areas associated with cognitive (PFC and hippocampal formation), motor (motor-sensory cortices), and sensory (audiovisual cortices) functions. **b**, Diagram showing the cosine similarity of frequency distribution of 13 three-node motifs for 50 brain regions. Four dashed areas correspond to PFC, hippocampal formation, motor-sensory cortices, and audiovisual cortices, respectively. **c**, Spatial distribution of neurons in the isocortex^14^ (left) and hippocampal regions shown in the folded Y-Z view^13^ (right). Each dot represents a soma from the single-neuron connectome dataset, colored according to the cluster assignment of its corresponding brain region. **d**, Schematic diagram illustrating differences in motif frequency among all areas shown in the four classes of brain areas (see b) for four motifs that showed significant differences between the four classes. The remaining nine motifs showed relatively small differences among brain region classes (see **Fig. S7** for more details). Here, * indicates *P* < 0.05, ** indicates *P* < 0.01, *** indicates *P* < 0.001, and **** indicates *P* < 0.0001. **e**, Representative example brain regions from the four major classes, including frontal pole cortex (FRP, *n* = 222 neurons), hippocampal CA1 region (*n* = 4396), primary motor cortex (MOp, *n* = 957), and primary auditory area (AUDp, *n* = 541), shown in the standard mouse brain template. **f**, Diagrams depicting the frequency distribution of various motifs in connectome data from FRP, CA1, MOp (see **Fig. S8** for more details), and AUDp brain regions.

A total of 34,572 neurons across these four clustered regions were measured in terms of their spatial distribution (**Fig. 2c**). We found that the PFC and hippocampus exhibited preferential enrichment of non-recurrent motifs 2, 4, and 8. In contrast, the motor-sensory and audiovisual cortices showed preferential enrichment of recurrent motif 7, whereas the motor-sensory cortices had the lowest levels of motif 8 (**Fig. 2d**). In addition, from these four clusters of brain regions, we selected four most representative regions (FRP, CA1, MOp, and AUDp) to visualize their single-neuron projection patterns (**Fig. 2e, see Fig. S1** and **S2** for more details), whereby the frequency distributions of the 13 connectivity motifs across these four regions also revealed substantial distributional differences (**Fig. 2f**).

We also found that these preferential motif distributions were stable and not constrained by the traditional stringent criterion of defining synaptic connections based on an axon-dendrite distance of < 5 µm. We subsequently tested different distance thresholds (3, 7, 9, and 11 µm; **Fig. S3**) and found that these variations did not significantly alter the motif distributions. Furthermore, we confirmed that the observed motif distributions were not driven by subsampling of neurons (20% to 100%, **Fig. S4**), the choice of specific regions (intratelencephalic IT region, **Fig. S5**), or pruning of synapses (0% to 40%, **Fig. S6**). This finding is important, as it indicates that motif distributions can serve as a metric for distinguishing connectivity patterns across different brain regions, independent of local microscopic statistics.

### Incorporating intracortical connectivity motifs in RNN and Transformer using CINA

Incorporation of connectivity architecture revealed by single-neuron connectomes into ANNs (represented by both RNN and Transformer) faces the challenge of converting key features of connectivity into equations compatible with ANN calculations. We developed an algorithm, referred to as “Connectome-Informed Neural network Algorithm” (CINA), based on the idea that recurrently connected ANNs could be constructed from 3-node connectivity motifs of various frequency distributions. Previous studies on counting motif frequency in the brain^12^ and computational models^15^ have used a serial sampling approach (**Fig. 3a**), which is computationally demanding due to the requirement of exhaustive serial motif search in large networks. Our CINA uses an efficient matrix multiplication-based method for simultaneous calculation of motif frequencies for all 13 three-node motifs (**Fig. 3b**). For the connectivity matrix, we first defined a method to obtain the frequency statistics of different motif categories within a small network of three neurons. This is achieved by performing element-wise probabilistic multiplication of matrix entries. Subsequently, we extend this approach to arbitrary neuronal networks using mathematical matrix operations, converting the element-wise multiplication into matrix multiplication. This key breakthrough allows the incorporation of the motif distribution into both RNNs and Transformers (here we use Mamba^16^ for example) based on combined use of a loss function and task error function (**Fig. 3c, d**). Both spiking and non-spiking neurons were integrated into these two types of architectures, and approximate gradient descent algorithm^17^ was employed for network training. During network learning, the CINA could gradually embed the topographic structure of biological networks, in the form of motif distribution, into the recurrent matrix weights of RNNs and Transformers (**Fig. 3e**). We further compared the computational time of our proposed matrix-based method against that of the traditional serial approach. The result showed that the speedup of our method increased with neuron number, reaching approximately 300-fold at 1,024 neurons (**Fig. 3f)**, demonstrating its practical utility.

**Fig. 3.**
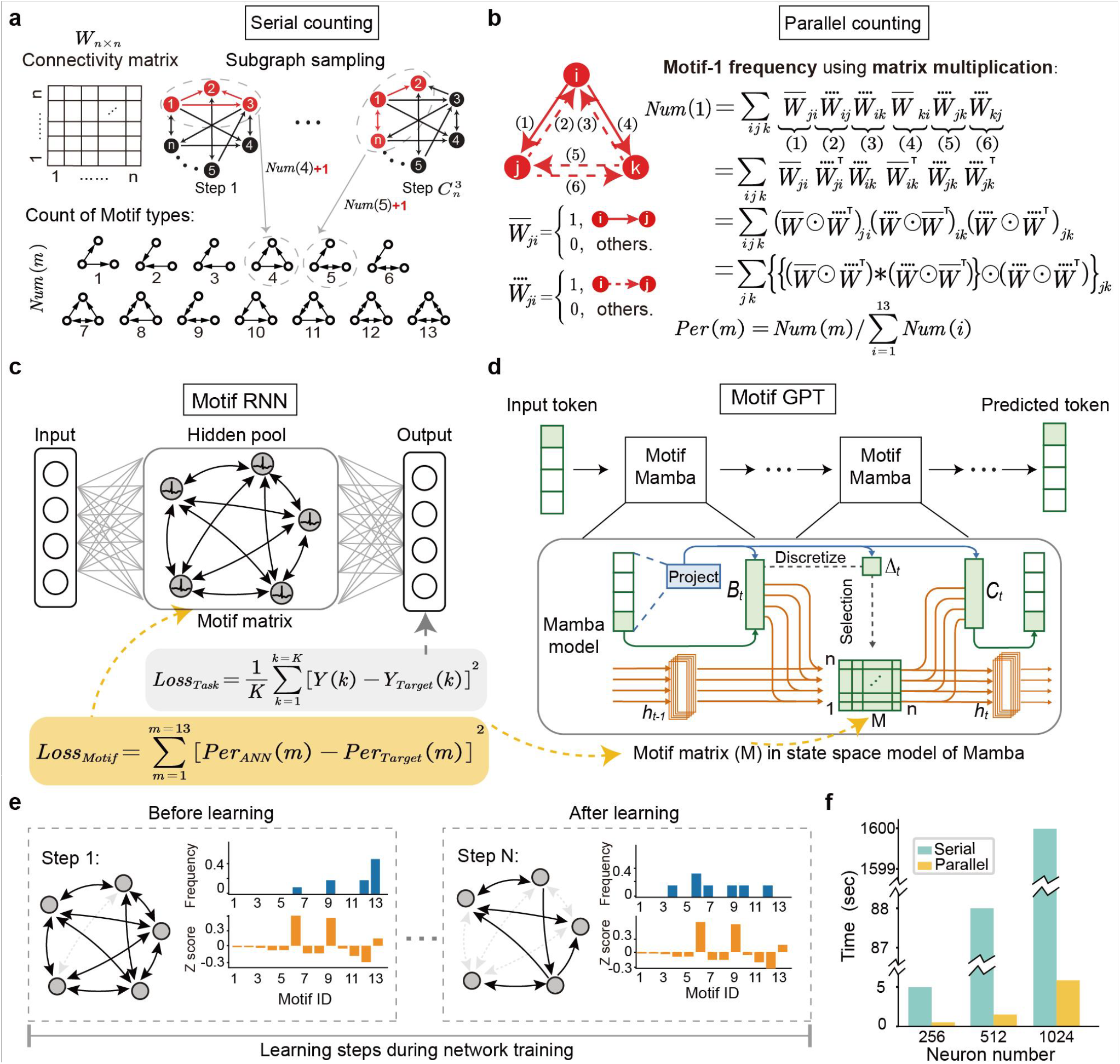
Connectome-informed neural network algorithm for embedding motif distributions. **a**, Schematic diagram depicting the serial motif counting method using sequential motif sampling and classification, requiring 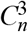 steps of sampling, each counting resulted in addition of number “1” into the total number of the specific motif [“*Num(m)*”]. Red node, neurons chosen for each step of motif counting. The 13 motifs (indexed 1 to 13) are shown below. **b**, Diagram depicting the CINA approach that calculates “*Num* (*m*)” through a parallel computation function using the adjacency matrix 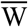 with Hadamard product (⊙) and matrix multiplication (*) operations (see Methods). **c**, Network architecture of Motif-RNN. The input and output layers are fully connected to a recurrently connected hidden pool of non-spiking (ANN) or spiking (SNN) neurons. The network motifs in the hidden pool were endowed by CINA via the “ *Loss*_*Motif*_ “ function (equation above, where “ *Per*_*ANN*_ (*m*) “ and “ *Per*_*Target*_ (*m)* “ represent the percentage of a specific motif in the ANN and target neural network, respectively) during network learning with “*Loss*_*Task*_ “ function (equation below, where “*Y*(*k*) “ and “*Y*_*Targer*_(*k*) “ denote the *k*^*t*h^ output of the ANN and the corresponding task target, respectively). **d**, Network architecture of Motif-Mamba. Input tokens are processed by stacked Motif-Mamba blocks to generate the predicted token. In the Mamba state-space model, the original state-transition matrix is replaced by a motif-compatible square matrix (*M*, Motif-matrix), allowing CINA to endow the model with biologically derived motif distributions while retaining the selective state-space components (*B*_*t*_, *C*_*t*_, and Δ_*t*_). **e**, Diagrams illustrating an example of a training process constrained by biological connectivity motif, the initial hidden pool configuration with full recurrent connections and progressive CINA-endowed motif configuration after the *N*^*t*h^ step of learning. **f**, Comparison of the time required for serial counting and parallel CINA counting of various motifs, for increasing number of neurons.

### CINA-endowed motif structures enhanced RNN’s learning capability

We next evaluated the performance of RNNs with CINA-endowed motif distributions in performing two categories of learning tasks. First, for noise-resistant categorization tasks, we examined the network learning of (1) visual perception using perceptual decision-making (PDM) dataset^18^, (2) static image recognition using N-MNIST dataset^19,20^, (3) dynamic video recognition using DVS-Gesture dataset^21^, and (4) spoken audio recognition using TIDIGITS dataset^22^ (**Fig. 4a**). Second, we examined motor learning of four benchmark tasks, using OpenAI MuJoCo^23^ paradigm of the reinforcement learning for continuous robot control, including (1) balancing a single inverted pendulum (“Inverted pendulum-v2”), (2) stabilizing a chaotic inverted double pendulum (“Inverted double pendulum-v2”), (3) teaching a robot to walk (“Walker-v2”), and (4) enabling an ant to run (“Ant-v2”) (**Fig. 4b**). We used CINA to endow five distinct frequency distributions of three-node motifs in ANNs as follows: (1) Averaged motif distribution obtained by averaging those found in FRP and MOp cortices (“FRP/MOp-averaged”), (2) cortex-specific motif distribution found in either FRP or MOp cortex (“FRP-specific” or “MOp-specific”), and (3) artificially increased bias of motif distribution by applying a non-linear transformation to normalized *Z* scores for FRP- or MOp-specific motif distribution (“FRP-enhanced” or “MOp-enhanced”, **Fig. 4c, d**). As a control, we also tested the performance of ANNs with random connectivity (“vanilla”).

**Fig. 4.**
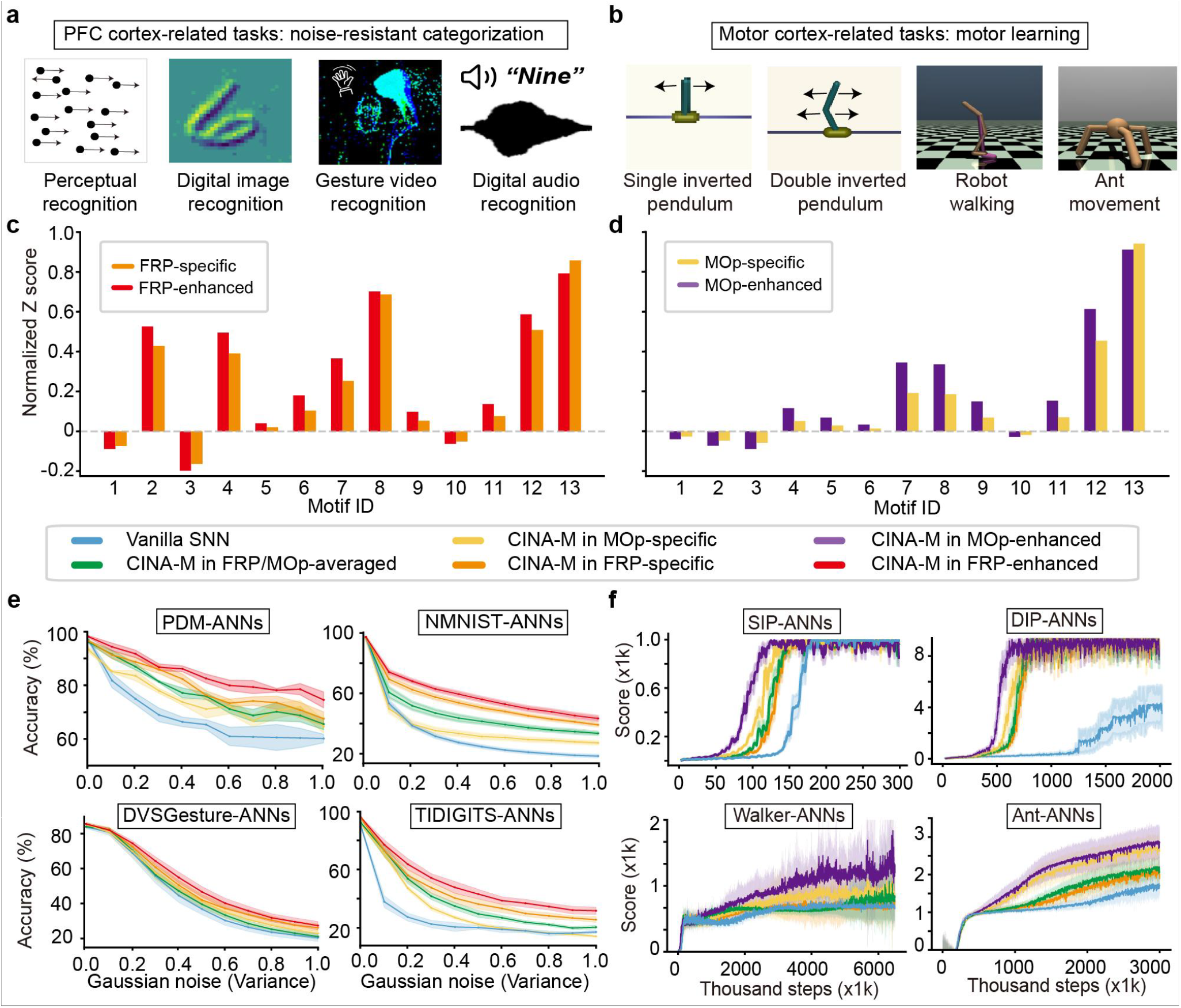
CINA-endowed biological and artificially biased motif distributions enhance task performance of RNNs. **a-b**, Images depicting various learning tasks examined for the performance achieved by RNNs endowed with various 3-node motif distributions. **a**, Four noise-resistant categorization tasks (PFC-related): perceptual decision-making (PDM), static image recognition (10-class digits recognition, N-MNIST, also abbreviated as NMNIST in figures, denotes the neuromorphic version of MNIST), temporal motion recognition (11-class video gestures, DVS Gesture), and voice recognition (10-class spoken digits, TIDIGITS). **b**, Four motor learning tasks (motor cortex-related) from OpenAI MuJoCo: balancing a single inverted pendulum (Inverted pendulum-v2, “SIP”), a chaotic inverted double pendulum (Inverted double pendulum-v2, “DIP”), teaching a robot to walk (Walker-v2), and enabling ant movement (Ant-v2). **c-d**, Diagram depicts the motif distributions of “FRP-specific”, “MOp-specific” and artificially biased versions (“FRP-enhanced”, “MOp-enhanced”). For “FRP-enhanced” and “MOp-enhanced” distributions, the original normalized *Z* scores (*NZ*) were first artificially biased (positive *NZ* scores were amplified using the function *ln* (1+5 *NZ*) and negative *NZ* scores were attenuated by mapping them as *ln* (1+3 ∣*NZ* ∣)), then renormalized. For visualization, all *NZ* were further transformed using *log*_10_(1+8*NZ*). **e-f**, Performance of CINA-endowed RNNs with the “vanilla” control in the four noise-resistant categorization tasks **e**, shown as accuracy under increasing input noise variance, and four motor learning tasks **f**, shown as reward score during reinforcement learning. Different motif distributions are color-coded (box above).

For the PDM task (**Fig. 4e**), we found that the mean accuracy (across all input noise levels) for RNNs with “FRP-specific”, “FRP/MOp-averaged”, and “MOp-specific” motif distributions were significantly higher than that found for the “vanilla” distribution by 6.6 ± 1.6% (SEM, *n* = 5 repetitions, *P* = 0.004), 5.2 ± 1.4% (*P* = 0.013), and 4.0 ± 1.9% (*P* = 0.038), respectively, indicating enhanced reliability of performance against input noise. Among these three distributions, “FRP-specific” distribution showed the highest accuracy (reliability) for the PDM task, in line with the notion that this task involves maintaining stable decision-making under noise, a known function of FRP^24^. Interestingly, “FRP-enhanced” distribution achieved accuracy surpassing that of the biologically relevant “FRP-specific” distribution by 2.5 ± 0.9% (*P* = 0.032). The results on the performance of CINA-endowed RNNs on three other noise-resistant categorization tasks (i.e., static clothing classification, dynamic video recognition, and spoken digit recognition tasks, **Fig. 4e**) were very similar to that described above for the PDM task.

For two single inverted pendulum and double pendulum motor learning tasks, we found that although the accuracies of task performance (as reflected by reinforcement reward scores) were similar (**Fig. 4f**), the learning speed (convergence rate) of ANNs with CINA-endowed “MOp-specific”, “FRP/MOp-averaged”, and “FRP-specific” motif distributions were significantly faster than that found for the “vanilla” distribution by 27.7 ± 1.8% (SEM, *n* = 20 repetitions, *P* < 0.001), 17.8 ± 1.3% (*P* < 0.001), and 16.6 ± 1.5% (*P* < 0.001) for inverted pendulum-v2 task, respectively; by 73.7 ± 2.0% (*P* < 0.001), 69.6 ± 2.2% (*P* < 0.001), and 64.0 ± 1.6% (*P* < 0.001) for inverted double pendulum-v2 task, respectively. Thus, “MOp-specific” distribution yielded the fastest learning speed on the pendulum tasks, in line with the notion that learning of these motor tasks were facilitated by motif distribution of the motor cortex. Moreover, “MOp-enhanced” motif distribution achieved a learning speed significantly surpassed that of the biologically relevant “MOp-specific” distribution (15.1 ± 2.5%, *P* < 0.001, inverted pendulum-v2; 10.0 ± 1.7%, *P* = 0.015, inverted double pendulum-v2). The results on the learning speed of two other motor learning tasks (i.e., robot walking and ant movement tasks) were similar to that described above. Furthermore, the accuracies of these motor learning, as indicated by the reward scores, were also progressively elevated for RNNs endowed with “FRP-specific”, “FRP/MOp average”, “MOp-specific” and “MOp-enhanced” motif distributions than that of the “vanilla” distribution (**Fig. 4f**). For robot walking and ant movement motor learning tasks, reward scores were progressively elevated for RNNs endowed with “FRP-specific”, “FRP/MOp-average”, “MOp-specific”, and “MOp-enhanced” motif distributions than that of the “vanilla” distribution by 19.7 ± 5.3% (SEM, n = 20 repetitions, P = 0.006), 21.5 ± 6.8% (P = 0.013), 27.7 ± 10.0% (P = 0.024), and 46.2 ± 12.4% (P = 0.007) for robot walking task, respectively; by 31.0 ± 9.9% (SEM, n = 20 repetitions, P = 0.017), 46.0 ± 16.4% (P = 0.027), 68.9 ± 9.6% (P < 0.001), and 81.6 ± 17.6% (P = 0.002) for ant movement task, respectively.

Taken together, these results showed that RNNs endowed with “FRP/MOp-averaged” motif distribution in general achieved higher performance in all tasks than “vanilla” RNNs. Furthermore, in line with the respective functions of FRP and MOp, RNNs endowed with “FRP-specific” and “MOp-specific” motif distribution resulted in higher performance for noise-resistant categorization and motor learning tasks, respectively. Similar results were found when the activation function of RNNs were replaced with spiking neurons (**Fig. S9**).

### Motif-informed Transformer further enhanced LLM’s prediction capability

We selected Mamba as the basic transformer block, and then modified its state-transition matrix from the original structured diagonal form to a square matrix (*M*, termed the Motif-matrix in **Fig. 3d**), in order to make motif-distribution constraints applicable (named Motif-Mamba). We then used CINA to endow three distinct biologically derived motif distributions in Motif-Mamba as follows: (1) an averaged motif distribution obtained by averaging those found in FRP and MOp cortices, denoted “Average-Motif-Mamba”; (2) the cortex-specific motif distribution found in FRP, denoted “FRP-Motif-Mamba”; and (3) the cortex-specific motif distribution found in MOp, denoted “MOp-Motif-Mamba”. We next evaluated the performance of a 130M-parameter Mamba LLM endowed with CINA-constrained motif priors on 6 popular downstream zero-shot evaluation language benchmarks, including LAMBADA task for broad-context word prediction^25^, HellaSwag task for commonsense situation completion^26^, PIQA task for physical commonsense reasoning^27^, ARC-E and ARC-C tasks for easy-level and challenge-level science question answering and reasoning^28^, respectively, and WinoGrande task for resolving ambiguous words in a sentence^29^ (**Fig. 5a**). Together, these benchmark tasks assess complementary aspects of language prediction, ranging from long-context prediction to commonsense and scientific reasoning.

**Fig. 5.**
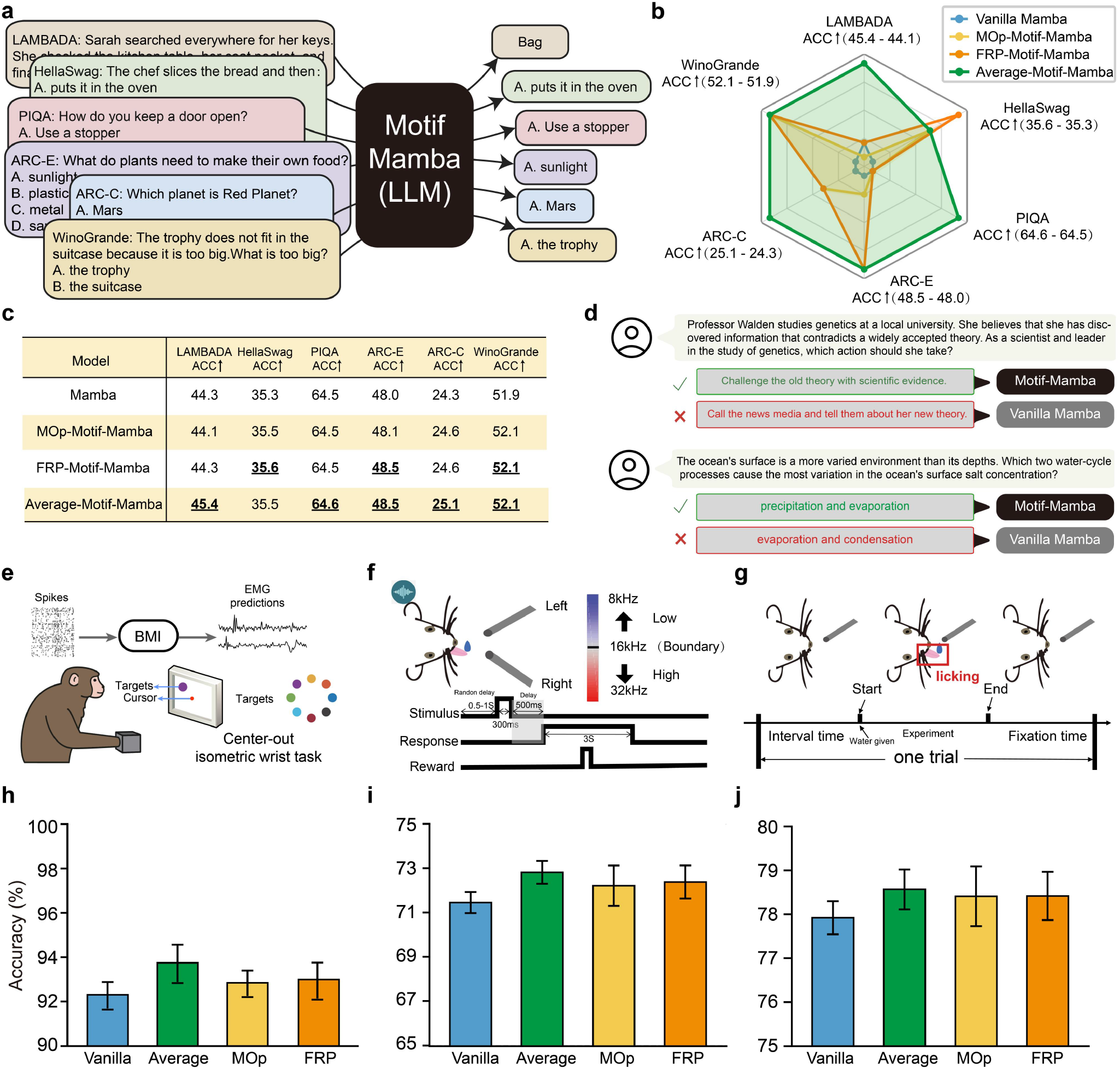
Biological motif priors embedded to Motif-Mamba on language benchmarks and brain-signal decoding tasks. **a**, Schematic diagram illustrating the natural language processing tasks. **b**, Radar plot showing accuracy on six downstream natural language question and answer benchmarks. For each benchmark, the values in parentheses indicate the maximum and minimum accuracies among the compared models. For visualization, the radial scale of each axis was set from 0.9 times the minimum accuracy to 1.1 times the maximum accuracy across the compared models for that benchmark. **c**, Accuracy values of the vanilla Mamba and motif-constrained Mamba models on the six downstream natural language question and answer benchmarks. Dashed and bold values indicate the best-performing model among the compared models for each benchmark. **d**, Representative examples from two natural language question and answer benchmarks, showing that Motif-Mamba correctly completes the tasks. **e**, Schematic diagram of the BMI decoding using center-out movement-direction classification task, in which invasive neural recordings were used to decode one of eight movement directions. **f**, Schematic diagram of the BMI decoding using mouse auditory two-alternative forced-choice task, in which animals classified tones as low or high frequency by licking the left or right water port. **g**, Schematic diagram of the BMI decoding using mouse fixed-interval licking task, in which electrophysiological spike recordings were used to decode binary lick/no-lick behavior. **h-j**, Classification accuracy of the vanilla Mamba and motif-constrained Mamba (FRP-Motif, MOp-Motif, and Average-Motif) models on the three brain-signal decoding tasks shown in e-g, respectively.

We found that relative to the vanilla Mamba, biologically derived motif priors produced additional gains (**Fig. 5b, c**), “MOp-Motif Mamba”, “FRP-Motif-Mamba”, and “Average-Motif-Mamba” remarkedly improved the average accuracy across the all 6 language benchmarks by 0.22%, 0.45%, and 1.12%, respectively. Among them, “Average-Motif-Mamba” gave the strongest overall performance (improving accuracy by 2.48%, LAMBADA; 0.57%, HellaSwag; 0.16%, PIQA; 1.04%, ARC-E; 3.29%, ARC-C; 0.39%, WinoGrande relative to the vanilla Mamba), supporting the previous hypothesis that the averaged cortical motif distribution might serve as a more generic and broadly transferable biological prior. In addition, “FRP-Motif Mamba” showed advantages on selected benchmarks (0.85%, HellaSwag; 1.04%, ARC-E; 1.23%, ARC-C; 0.39%, WinoGrande) relative to the vanilla Mamba, whereas “MOp-Motif-Mamba” produced only limited improvement in this setting. Representative examples illustrated two benchmark questions that were correctly answered by Motif-Mamba (**Fig. 5d**). A plausible interpretation is that these language benchmarks are better matched to the cognitive cortical prior represented by the FRP region than that by the Mop region (motor-related prior).

In addition to language benchmarks, we further found that CINA-constrained motif priors could improve brain-signal decoding tasks related to BMI applications. We selected three neural decoding benchmarks: (1) a center-out classification task for decoding one of eight movement directions from invasive neural recordings^30^ (**Fig. 5e**); (2) a mouse auditory two-alternative forced-choice task for decoding low- or high-frequency tone choices by licking the left or right water port^31^ (**Fig. 5f**); (3) a mouse licking task for decoding binary lick/no-lick behavior from electrophysiological spike recordings^32^ (**Fig. 5g**). Experimental results indicated that, relative to the vanilla Mamba, biologically derived motif priors produced additional gains on all these three tasks (up to 93.7%, 72.8%, 78.6%, **Fig. 5h, i, j**), where “Average-Motif-Mamba” gave the strongest overall performance, improving classification accuracy by 1.56 ± 0.36% (SEM, n = 5 repetitions, *P* = 0.012), 1.91 ± 0.34% (*P* = 0.006), and 0.83 ± 0.11% (*P* = 0.040) for the center-out movement-direction classification task, mouse auditory forced-choice and mouse spike-to-behavior prediction tasks, respectively.

Together, these results demonstrated that our method achieved higher accuracy simultaneously in both language-based prediction and brain-signal decoding tasks, suggesting that connectome-derived motif priors can provide additional structural knowledge to LLM.

### Topological organization of hidden pool neurons in specific motif-endowed ANNs

To understand the network properties in ANNs endowed with various motif distributions from the perspective of the graph theory, we systematically analyzed network structure of hidden pool neurons by quantifying the topological structures of post-learning connectivity weights and the distribution of their connectivity. Complex networks often naturally comprise distinct communities or modules^33^. We first examined whether embedding motif distributions into RNNs could drive the network connectivity towards modular structures similar to real-world networks. We employed the Infomap algorithm^34,35^ to detect the community structures of RNNs endowed with different motif distributions, using optimal modularity as the criterion for community partitioning. The community partitions for various RNNs are shown in **Fig. 6a**, which displays at most top 15 strongest communities for “FRP-specific”, “MOp-specific”, and “FRP/MOp-averaged” motif distributions, and each had distinct community structures. These structures were all distinct from that found for the “vanilla” RNN, which exhibits no discernible community. We calculated modularity using the directed modularity method^36^, quantified by the modularity index, and compared it against the baseline of ER random networks^37^ with matched graph size and sparsity. The RNNs endowed with “FRP-specific”, “MOp-specific”, and “FRP/MOp-averaged” motif distributions exhibited markedly higher modularity indices than the “vanilla” RNN, with the “MOp-specific” motif distribution yielding the highest index (**Fig. 6b**).

**Fig. 6.**
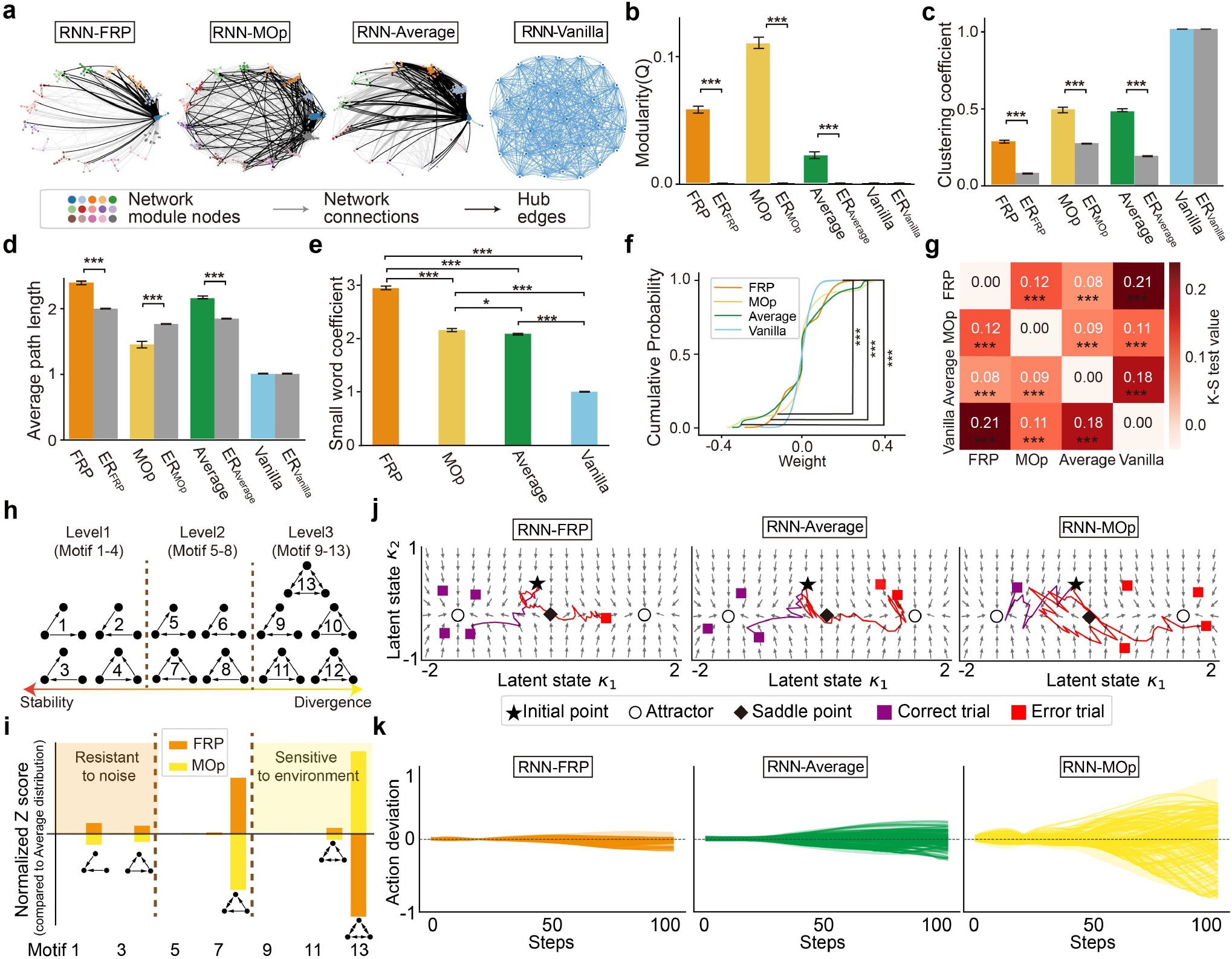
Motif-constrained RNNs exhibit distinct topological properties and latent dynamics. **a**, Network topological structures after community partitioning of RNNs endowed with biological motif distributions and the “vanilla” distribution. Different colors represent distinct communities, with intra-community connections shown in the corresponding community color, inter-community connections shown in gray, and the top 20% most important connections (measured by node degree) highlighted in black. We sampled 25 nodes from the total of 512 to more effectively visualize the network structure of the “vanilla” RNN, which does not exhibit a clear community structure. **b-e**, Diagrams depicting network modularity index (b), clustering coefficient (c), averaged path length (d) and small-world coefficient (e). For each motif distribution, 10 RNN repetitions models were trained and compared against 100 ER random networks. Here, * indicates P < 0.05 and *** indicates P < 0.001. **f**, The cumulative distribution curves of connectivity weights for RNNs endowed with different motif distributions and the “vanilla” RNN. **g**, The heatmap depicts the statistic value *D* of the Kolmogorov-Smirnov test for each pair of RNNs and *P* value, where *D* > 0.0054 denoting *P* < 0.001. The number of observations corresponds to their connectivity elements. **h**, Diagram of all 13 three-node motifs organized into a three-level hierarchy based on their local dynamical behavior. **i**, Normalized *Z* scores of the “FRP-specific” and “MOp-specific” motif distributions relative to the “FRP/MOp-averaged” distribution, mapped onto the hierarchy in **h. j**, Latent-state dynamical trajectories during sample trials of the PDM task for RNNs endowed with biological motif distributions. Collective activity was projected onto a two-dimensional latent space by low-rank method^41^. **k**, Diagrams depicting statistical tracking of action deviation across recurrent steps for 200 normalized observations in the corresponding RNNs during the SIP task. Each line represents one trial. The *x* axis indicates recurrent iteration steps and the *y* axis indicates action deviation.

The small-world property is often a feature of complex networks that reflects the balance between local clustering and global integration, supporting efficient information transmission^38,39^. The small-world coefficient is used to assess whether a network has an efficient organizational structure and is quantified using two key parameters: the clustering coefficient (**Fig. 6c**) and the average path length (**Fig. 6d**; see Methods). Our results showed that RNNs endowed with different distributions of connectivity motifs displayed distinct differences in these two parameters among themselves and from those of the “vanilla” RNN (**Fig. 6c, d**), leading to significant small-world coefficients in all RNNs endowed with biological motif distributions (**Fig. 6e**). Notably, RNNs endowed with “FRP-specific” motif distribution exhibited the highest small-world index, suggesting that this motif structure represents a more optimized balance between local clustering and global transmission. We further constructed weight matrices of hidden pool neurons for RNNs endowed with different motif distributions (**Fig. S10**) and found that RNNs endowed with three types of biologically inspired motif distributions showed sparse and distinct patterns of weight distributions (**Fig. 6f**), whereas the RNN with the “vanilla” distribution exhibited random weights. The statistically significant differences were further validated by the cumulative probability analysis of the weight distributions (Kolmogorov-Smirnov test^40^: *D* > 0.11, *P* < 0.001) (**Fig. 6g**) and among themselves (*D* > 0.08, *P* < 0.001). Notably, the greatest difference was found between the “vanilla” RNN and RNN with CINA-endowed “FRP-specific” motif distribution (*D* = 0.21, *P* < 0.001).

To further examine the dynamical basis of the functional differences conferred by cortex-specific motif distributions, we analyzed the internal dynamics of the trained recurrent networks. We found that, based on their stability and divergence, the 13 three-node motifs could be organized into a three-level hierarchy (**Fig. 6h**): Level one consists of motifs 1, 2, 3, and 4; Level two consists of motifs 5, 6, 7, and 8; and Level three consists of motifs 9, 10, 11, 12 and 13. This hierarchical organization suggests that the functional capabilities of different brain regions can be inferred by analyzing the enrichment of motifs from different levels within each brain region. Our analysis confirmed this hypothesis. For example, relative to the “FRP/MOp-averaged” profile, the “FRP-specific” motif distribution was shifted toward the more stable side of the hierarchy, suggesting that the FRP may play a role in tasks requiring network stability. Conversely, the “MOp-specific” distribution was shifted toward the more recurrent side, indicating that the MOp may be more involved in reinforcement learning-type exploratory tasks (**Fig. 6i**). Supporting this, latent-state dynamics analysis during the noise-resistant PDM task showed that RNNs endowed with the “FRP-specific” motif distribution achieved the highest accuracy and the most robust convergence toward stable decision points under noise (**Fig. 6j**).

In addition, to assess how biological motif distributions shape action-level dynamics during motor control, we tracked action deviation across 200 normalized observations in the SIP task and found that the RNN endowed with “MOp-specific” motif distribution exhibited the broadest range of action deviation across recurrent steps (**Fig. 6k**). Together, these results suggest that cortex-specific motif distributions bias collective network dynamics toward distinct computational modes, suggesting that “FRP-specific” and “MOp-specific” profiles favoring noise-resistant and environment-sensitive computation, respectively.

## DISCUSSION

Single-neuron connectome data provide neuronal level information on the brain’s organizational principles that could help to design more efficient and capable ANN architectures. Using the available mouse intracortical connectome data, we identified the frequency distributions for all 13 possible 3-node connectivity motifs across 50 brain regions and developed a computational algorithm CINA to endow various motif distributions into the hidden-pool connectivity of ANNs. To further investigate functional consequences of such motif organization in ANNs, we selected frontal pole of PFC (FRP) and primary motor cortex (MOp) as two representative cortical areas involved in perceptual and motor functions, respectively. We found that ANNs with “FRP-enhanced” and “MOp-enhanced” motif distributions achieved the highest accuracy and efficiency in noise-resistant categorization and motor learning tasks, respectively. This is consistent with the crucial function of PFC in maintaining stable decision-making in the presence of noise interference^24^ and MOp is essential for movement planning and execution^42^. Further graph theory-based studies of network topography showed that ANNs endowed with FRP-specific, MOp-specific, or “FRP/MOp-averaged” motif distributions showed strong modularity and small-world features that were absent in the “vanilla” ANN with random connectivity. These results support the notion that motif-endowed network architecture underlies the elevated efficiency of ANNs.

Our motif analysis of the mouse connectome showed that FRP and MOp cortices contain higher proportions of non-recurrent and recurrent connectivity motifs, respectively. Such cortex-specific motif distributions are likely to serve for distinct functions of information processing. Compared with MOp, FRP-enriched motif 2 provides convergent connections that contribute to information integration. In contrast, motif 13 recurrent connections were enriched in MOp, the motor area for mapping antecedent conditions (e.g., sensory cues and recent choices) onto upcoming actions^42^, where recurrent connections could promote rapid neuronal synchronization and persistent activity. These observations are in line with the previous finding that non-recurrent sparse connectivity reduces noise interference and enhances the reliability in categorization tasks^9^, whereas relatively high recurrent connectivity motifs in MOp may support the exploratory flexibility required for motor learning. Such cortex-specific motif organization may emerge during evolution and development for performing specific tasks of various cortices.

In this study, we found that artificially increasing the bias of motif distributions in FRP and MOp further enhanced the ANN’s performance (including both RNN and Transformer) in tasks associated with these cortical regions. These results provide additional evidence that specific connectivity motif profiles contribute to the learning of distinct computational functions. Moreover, the elevated performance of ANNs with artificially enhanced motif distributions also exemplifies biologically inspired optimization of ANN architecture for specific tasks.

Taken together, our work characterized the motif organization of various brain regions, developed a computational algorithm for incorporating biological motif structures into ANNs, and demonstrated the high performance of these ANNs in noise-resistant categorization and motor learning tasks. These findings highlight the potential of connectome-inspired network design for improving artificial neural systems while providing new insights into the functional significance of specific motif profiles in various brain regions.

## METHODS

### Construction of intracortical connectivity matrix

To measure the motif distribution of intracortical connections in the mouse cortex, we generated a neuron connection matrix based on the single-neuron mouse connectome data in the Digital Brain Database (https://www.digital-brain.cn/). These single-neuron projectomes were obtained by the fluorescence Micro-Optical Sectioning Tomography (fMOST) method on mouse brains that were virally labeled with the fluorescent protein to achieve sparse labeling of intracortical pyramidal projection neurons^43,44^, with a spatial resolution of 0.3×0.3×0.3 *μm*. The acquired single-neuron images (including soma, dendritic and axonal arbors) were traced and registered to the standard mouse brain template (i.e. the Allen CCFv3, ref.^13^) in order to obtain the spatial coordinates of the neuronal soma and axon/dendrite arbors (**Fig. 1a**). Single-neuron images from each intracortical region were projected simultaneously to the same template to determine connectivity between each pair of neurons.

For each putative synaptic contact from source to target neurons, we computed the closest Euclidean distance (ED) from every axonal arbor to dendritic arbor. Corresponding to the typical length of dendritic spines and the size of axonal boutons^45,46^, we designated the axon-dendrite arbor pair with ED < 5 *μm* (other ED thresholds are also considered in **Fig. S3**) as putative sites for synaptic contacts (**Fig. 1b**). The total number of putative contact sites was used to indicate the connectivity score from neuron A to neuron B (**Fig. 1c**). To estimate the connectivity score of a given neuron pair, we used the following equation^46^:

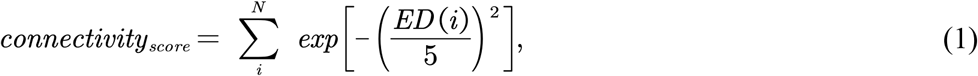

where *ED*(*i*) is the Euclidean distance for the *i*^*th*^ putative pre- and postsynaptic sites.

### Calculation of three-node motif distribution

We extended a previous matrix multiplication method^47^ for two-edge motif distribution calculation to all 13 possible three-node motifs. Let 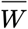 be the adjacency matrix of the connection matrix *W*, where 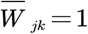 represents the presence of a directed connection from node to node *j*,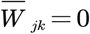 represents its absence. In contrast, to carve out unconnected edges, we define 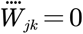 and 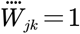 as the presence and absence of a directed connection from node from node *j* to node *k*, respectively. We define the vector where all elements are 1 as *L*=**1**_*N*,1_. For convenience, the number of 13 three-node motifs is calculated by the equation in a general form as follows:

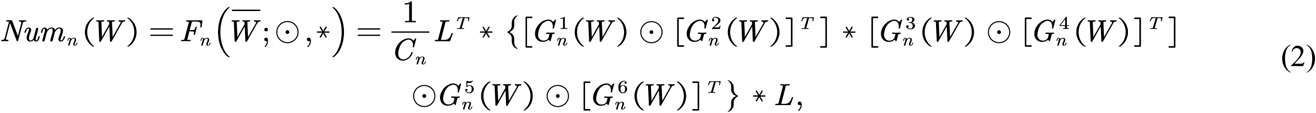

where ⊙ is the Hadamard product and * is the matrix multiplication, *n* is the ID number of three-node motifs, *C*_*n*_ is a constant. 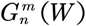 is mapped to be 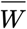 and 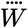, when the connection of the *m*^*th*^ edge of *n*^*th*^ motif is present and absent, respectively (see **Table S2** and **Table S3** for more details). The frequency of the *n*^*th*^ three-node motif is calculated by dividing its count by the total number of all three-node motifs. Note that the backpropagation algorithm was used to update model parameters by calculating the gradient of the loss function^17^. Since this calculation relies on the continuity and differentiability of the loss function, we approximate the binary adjacency matrix 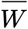 by setting 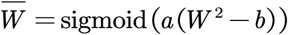, where *a* and *b* are adjustable parameters (**Fig. S11**).

The frequency of the *i*^*th*^ motif that differed from the frequency found in a randomly connected network is evaluated by calculating the *Z* score as utilized previously, *Z*_*i*_=(*Nreal*_*i*_ − *E*[*Nrand*_*i*_])/*STD* (*Nrand*_*i*_), where is the total number of the motif for the real network, *E*[*Nrand*_*i*_] and *STD* (*Nrand*_*i*_) are the mean and standard deviation of its total number of 100 ER random networks. Moreover, *Z* scores were normalized to unit vectors 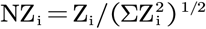 to enable direct comparison across different networks. To improve visualization of small normalized *Z* scores, we applied a nonlinear transformation *log*_10_(1+8*NZ*). This transformation preserves the sign and relative ordering of normalized *Z* scores, and all normalized *Z* scores. All normalized *Z* scores presented in figures throughout the manuscript refer to these transformed values unless explicitly stated otherwise.

### Analyze the stability of motif dynamics

For a given motif class *m*, the internal states **h**=[*h*_1_,*h*_2_,*h*_3_]^*T*^evolve according to

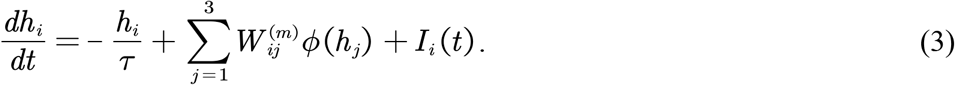

where *W*^*m*^ denotes the motif-specific recurrent coupling matrix *I*_*i*_(*t*), represents the external input drive. Following the Hartman–Grobman theorem, we characterized the local behavior of the nonlinear motif dynamics by linearizing the system around an equilibrium point **h**^*^. The Jacobian matrix **J** of the motif dynamics with respect to the state is **h** given by

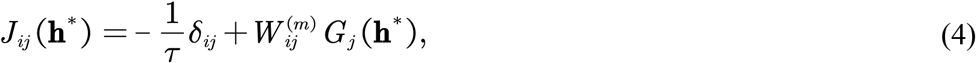

where *δ*_*ij*_ is the Kronecker delta and 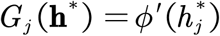 is the local gain at equilibrium. We then analyzed the eigenvalue conditions of the motif-specific Jacobian to derive sufficient stability conditions for different motif classes.

Based on these sufficient conditions, the 13 motifs were grouped into three dynamical levels. Level-1 motifs (IDs 1–4) form the most stable class and require only negative self-connections on all three nodes. Level-2 motifs (IDs 5–8) additionally contain a single reciprocal pair and require both negative self-connections and an extra stability constraint on that pair. Level-3 motifs (IDs 9–13) contain multiple reciprocal interactions or three-node cycles and require more stringent higher-order stability conditions. We therefore used this three-level grouping as a dynamical descriptor of motif composition in the present study.

This grouping was not intended to imply that isolated three-node motifs uniquely determine the full-network dynamics. Rather, it provided an interpretable local structural bias for comparing biological motif profiles. In this view, motif distributions enriched in Level-1 motifs were expected to favor more convergent and robust dynamics, whereas motif distributions enriched in Level-3 motifs were expected to favor more flexible and feedback-sensitive dynamics.

### Similarity analysis of motif distributions

To visualize the similarity of motif distributions across 50 brain regions, we performed hierarchical clustering on the 13-dimensional normalized *Z* score vectors of motif distributions (**Fig. 2a**). We calculated the cosine similarity between motif distribution vectors of different brain regions in **Fig. 2b** using the formula: 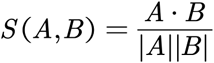, where *A* and *B* are the normalized *Z* score vectors of motif distributions for two brain regions. Moreover, we used Uniform Manifold Approximation and Projection^48^ (UMAP) to reduce the motif distributions of different brain regions to 2-dimensional embeddings, followed by K-means clustering to quantify the similarity among brain regions.

### ANN architecture for categorization and motor learning tasks

We used ANNs with an input layer, a hidden pool of recurrently connected neurons, and an output layer for visual/auditory categorization tasks or OpenAI MuJoCo^23^ reinforcement motor learning tasks. Network parameters are listed in **Table S4**. Audio data from TIDIGITS^22^ were preprocessed using Mel-frequency cepstral coefficients (MFCC) to transform 1D raw data into 2D frequency spectrum.

### Analysis of the modularity and small-world properties of ANNs

Community detection was performed using the Infomap algorithm^34,35^, and the quality of the community partition was evaluated by the optimal modularity, computed using Newman’s directed modularity^36^:

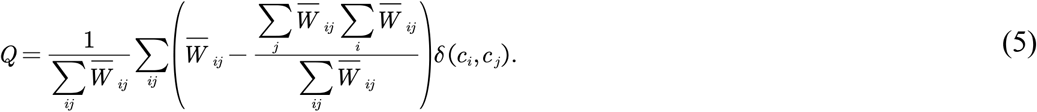

Here, 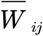 represents the existence of an edge from node *i* to node *j, c*_*i*_ denotes the corresponding community label for node *i*. *δ*(*c*_*i*_,*c*_*j*_)=1 if *i* and *j* belong to the same community, and 0 otherwise. The community-partitioned network is visualized using the Python package NetworkX^49^.

The Humphries–Gurney index^50^ is used to compute the small-world property, using the formula 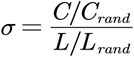 Here, is the clustering coefficient, is the average path length, *C*_*rand*_ and *L*_*rand*_ are the clustering coefficient and average path length of ER random networks, respectively. The clustering coefficient and average path length are computed as 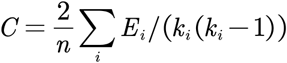 and 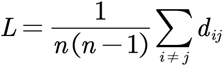, where *n* is the number of nodes, *E*_*i*_ is the number of edges among the neighbors of node *i, k*_*i*_, is the number of neighbors of node *i*, and *d*_*ij*_ is the shortest path length from node *i* to node *j*.

## Data and code availability

All data needed to evaluate the conclusions in the paper are present in the main text with citations. All source codes can be downloaded from https://github.com/I4m4I/motif.

## SUPPLEMENTARY MATERIALS

The list of supplementary materials for this article is shown as follows:

## Acknowledgements

This work was supported by the Brain Science and Brain-like Intelligence Technology - National Science and Technology Major Project (2025ZD0217200), Strategic Priority Research Program of the Chinese Academy of Sciences (Grant No. XDB1010302), CAS Project for Young Scientists in Basic Research (YSBR-116), Youth Innovation Promotion Association CAS, the Lingang Laboratory Fund (Grant No. LG-GG-202402-06-07, LGL-1987-09), the Shanghai Municipal Science and Technology Project (Grant No. 25ZR1401370, 25LN3200400), Special Support Project of Guangdong Province (Grant No.0720240209). The numerical calculations in this study were carried out on the ORISE Supercomputer.

## Author contributions

T.Z. designed the study. T.Z., Y.S., W.Y., J.Z., W.S., X.Z., C.H., X.Y.C., S.Z., S.J., Y.Y., X.C., X.X., M-m. P, Y.G.S., and B.X. performed the discussion and wrote the paper together. Y.S. contributed to the design of the theoretical framework. W.Y., J.Z., X.Z., C.H., X.Y.C., S.Z., and Y.Y. performed the experiments. S.J. contributed to the figures. W.S., X.C., X.X., and Y.G.S. contributed to the preparation of biological data.

## Competing interests

The authors declare no competing interests.

Supplementary Materials

## SUPPLEMENTARY FIGURES

**Fig. S1.**
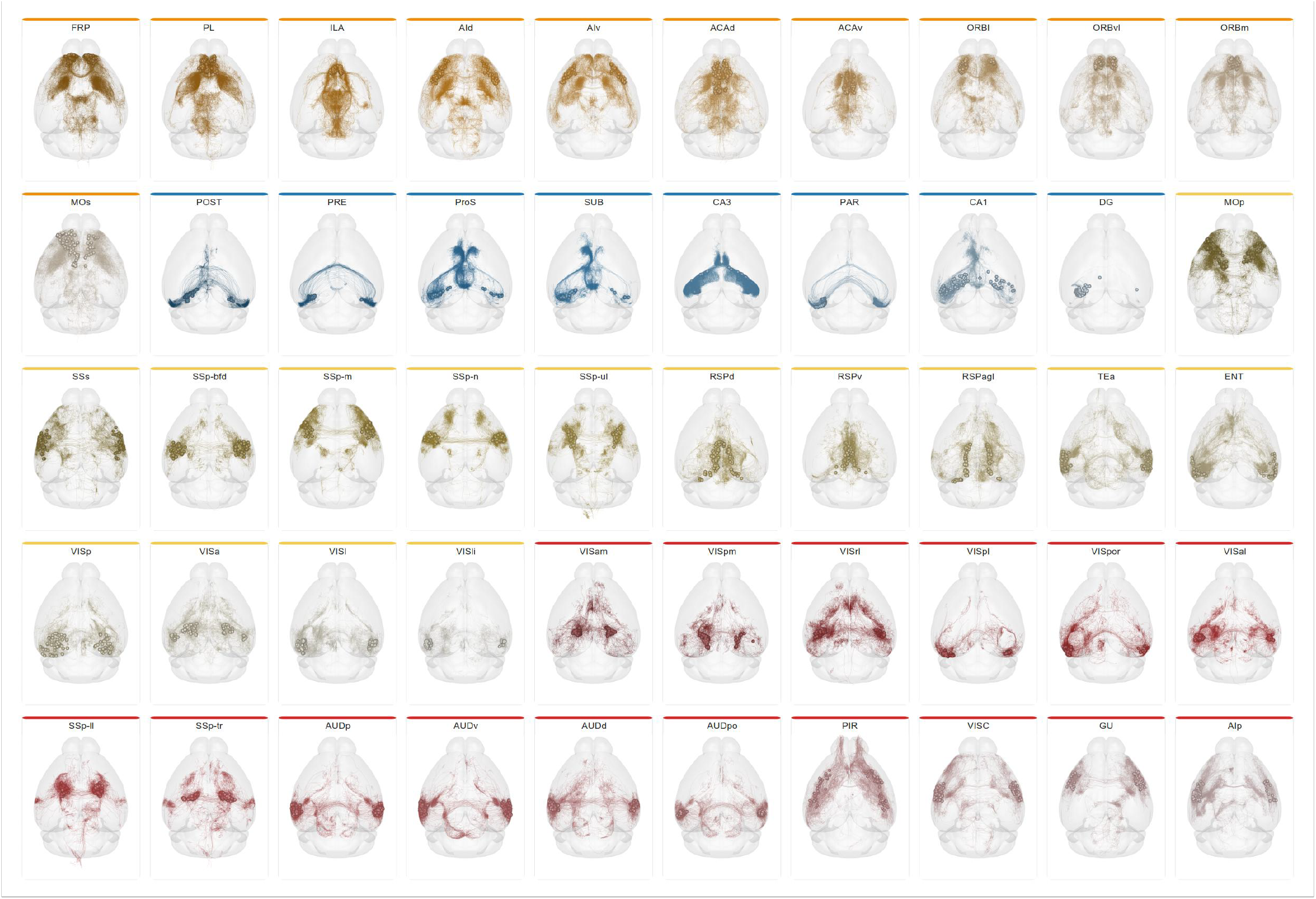
Spatial distributions of reconstructed neurons across cortical and hippocampal regions. Spatial distributions of reconstructed neurons in individual regions belonging to prefrontal cortex, hippocampal formation, motor-sensory cortices, and audiovisual cortices. Each panel shows neurons from one subregion projected onto a transparent whole-brain template, with colors indicating the corresponding brain region class.

**Fig. S2.**
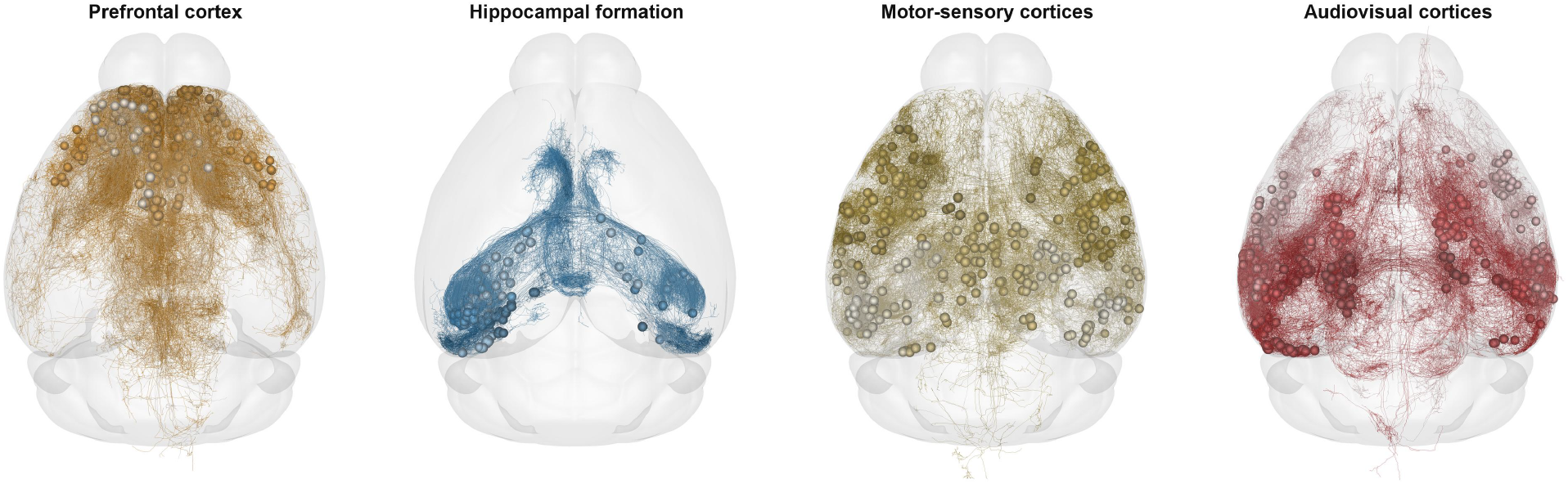
Grouped spatial distributions of reconstructed neurons across four region classes. Spatial distributions of reconstructed neurons grouped by prefrontal cortex, hippocampal formation, motor-sensory cortices, and audiovisual cortices. Each panel shows neurons from all included regions within one class, projected onto a transparent whole-brain template.

**Fig. S3.**
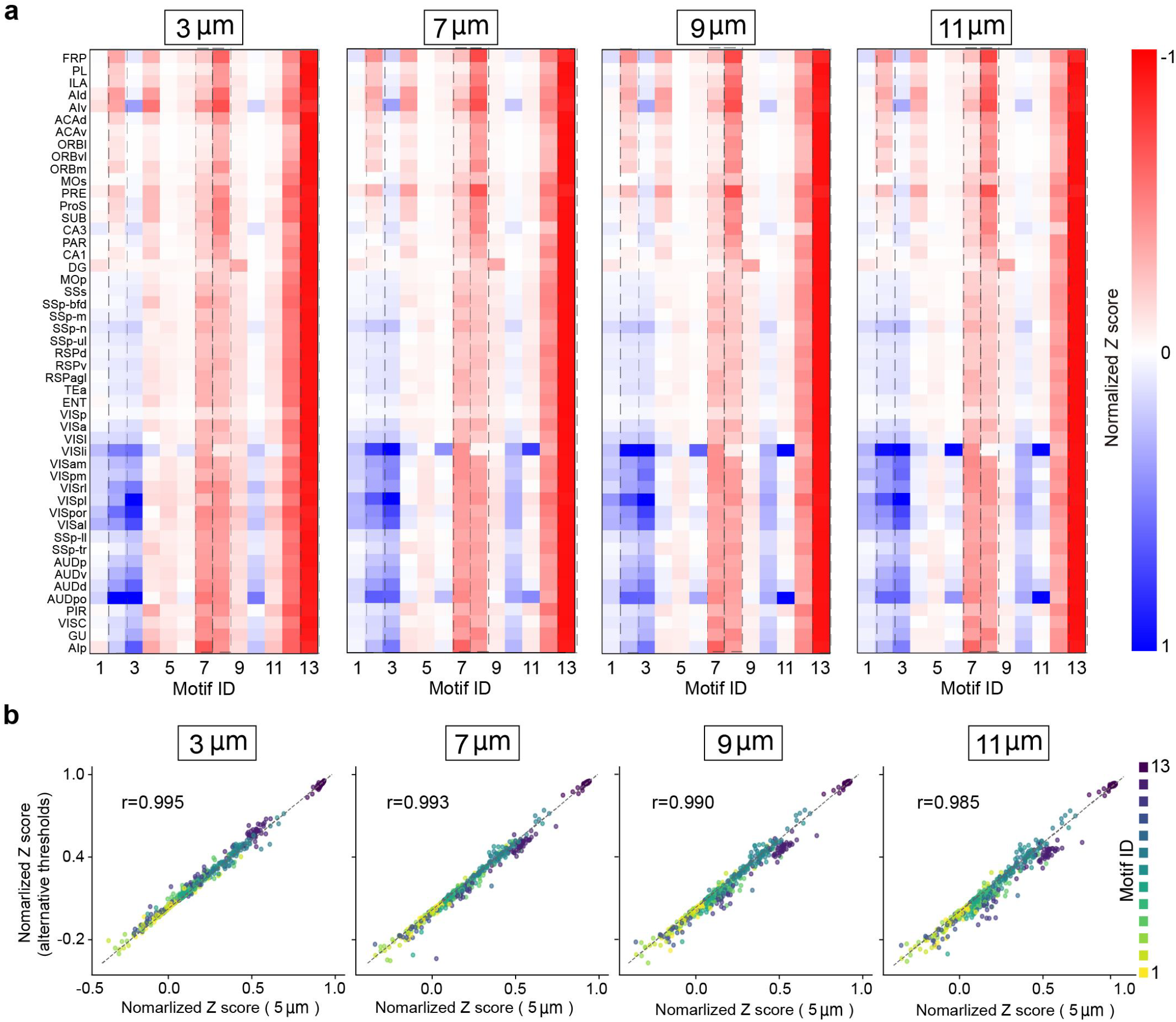
Similarity of motif distributions for various Euclidean distance (ED) thresholds. **a**, Diagram showing the similarity of the frequency distribution of 13 three-node motifs for 50 brain regions across various Euclidean distance (ED) thresholds. **b**, Schematic diagram illustrating the Pearson correlation of motif distributions between the baseline threshold (5 μ*m*) and alternative distance thresholds (3, 7, 9, and 11 μ*m*).

**Fig. S4.**
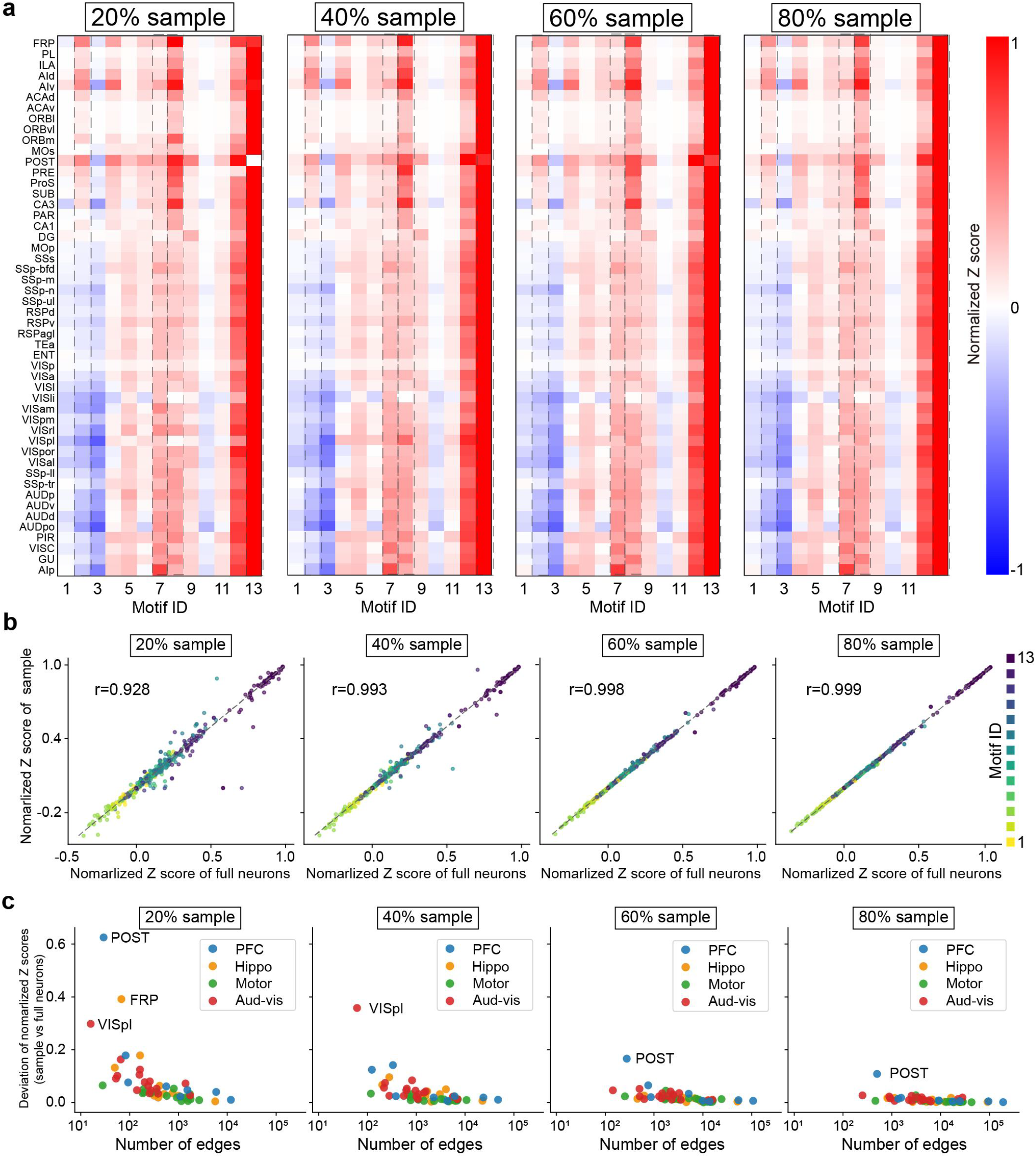
Similarity of motif distributions for various subsampling of neurons. **a**, Diagram showing the similarity of the frequency distribution of 13 three-node motifs for 50 brain regions across various pruning ratio of weak connection. **b**, Schematic diagram illustrating the Pearson correlation for 50 brain regions between random resampling of subsets (from 20% to 100%) of total neurons. **c**, Deviation of normalized Z-scores between subsampled and full neurons as a function of edge count across different brain regions.

**Fig. S5.**
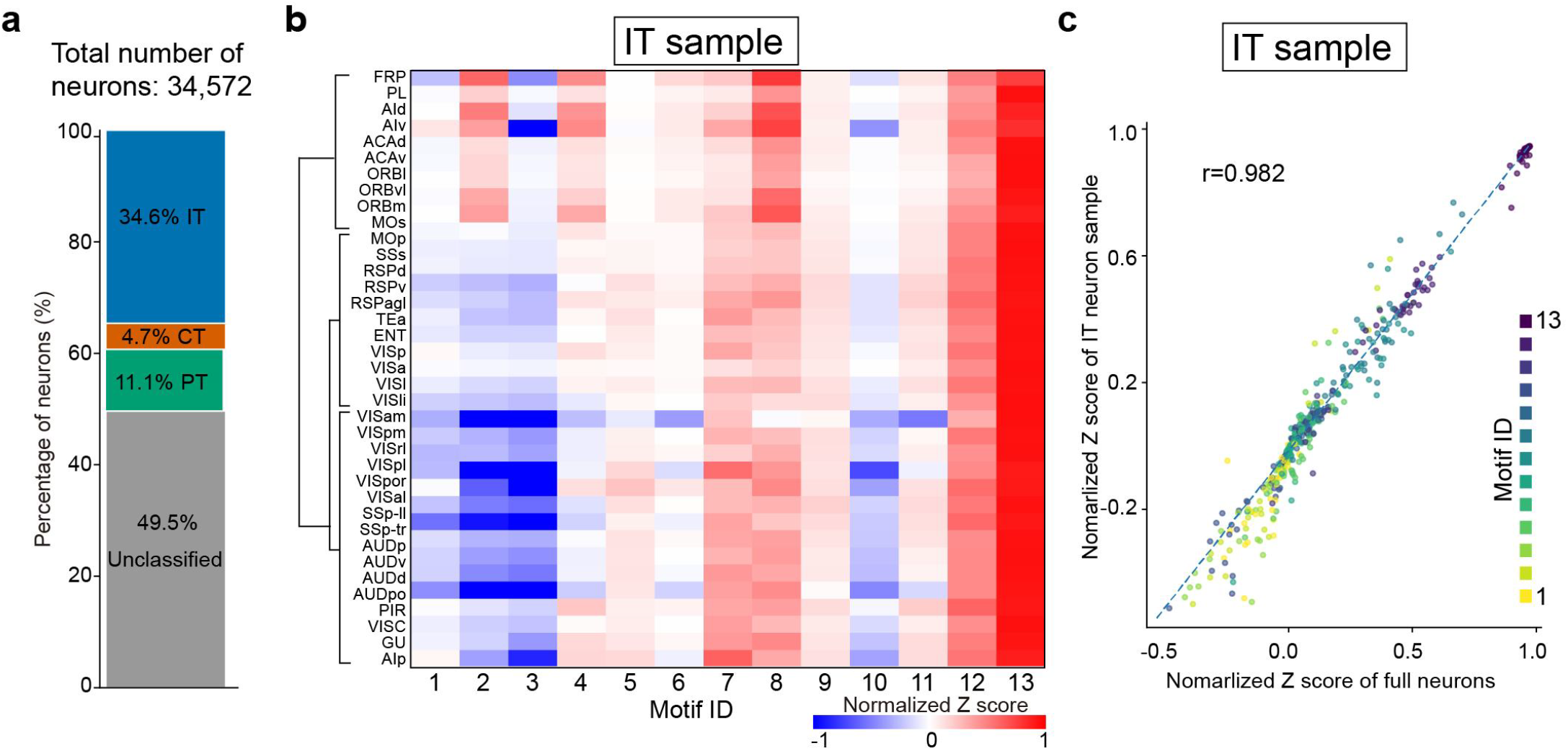
Motif distributions for various subsampling of neurons. **a**, The percentage of different type neurons in the total number neurons. Here, IT, PT, and CT are intratelencephalic, pyramidal tract, and corticothalamic neurons. **b**, The frequency distributions of 13 three-node motifs for 50 brain regions of IT neurons. **c**, Schematic diagram illustrating the Pearson correlation for 50 brain regions between IT sampling and the total number of neurons.

**Fig. S6.**
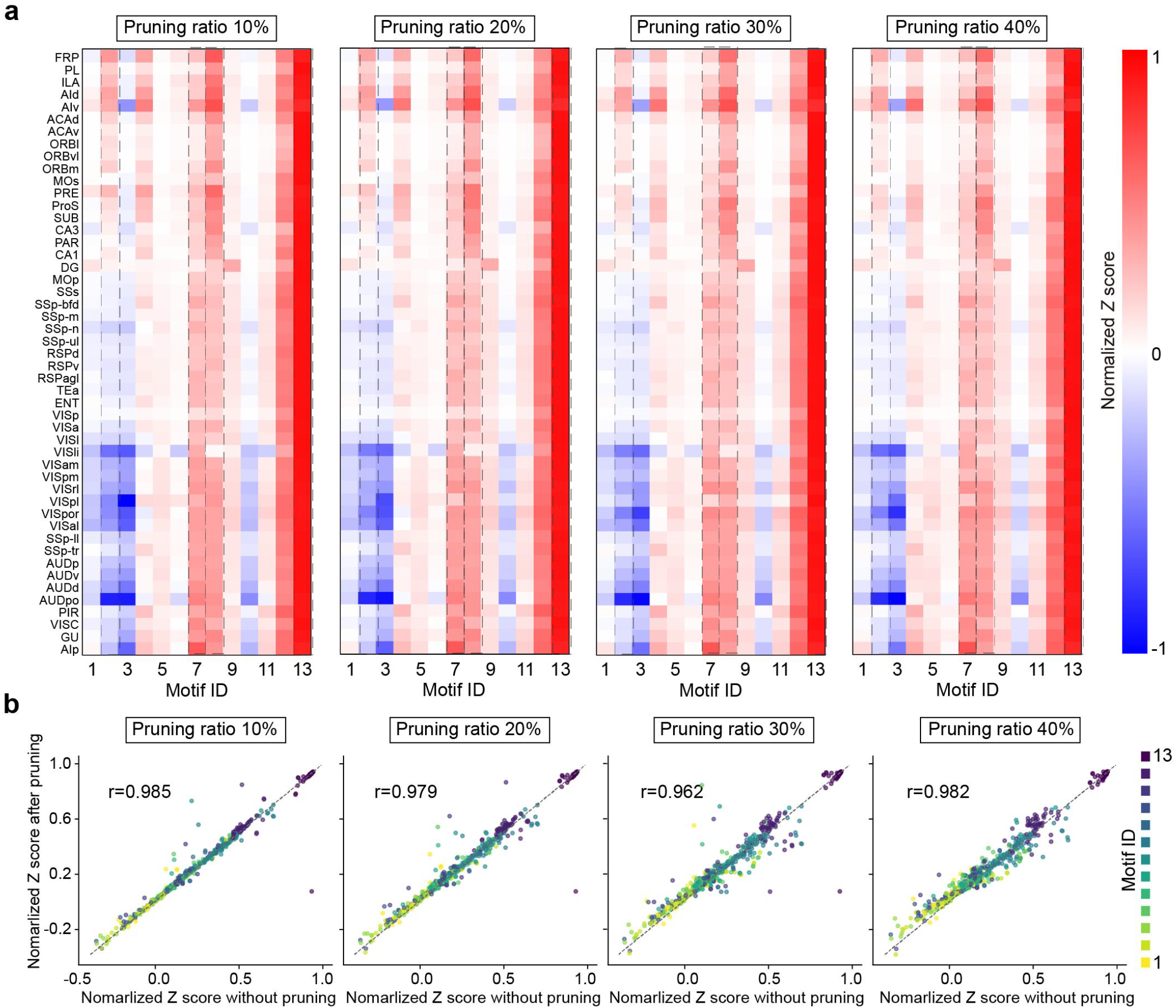
Similarity of motif distributions for various pruning ratios of weak connections. **a**, Diagram showing the similarity of the frequency distribution of 13 three-node motifs for 50 brain regions across various pruning ratio of weak connection. **b**, Schematic diagram illustrating the Pearson correlation for 50 brain regions across various pruning ratio of weak connection.

**Fig. S7.**
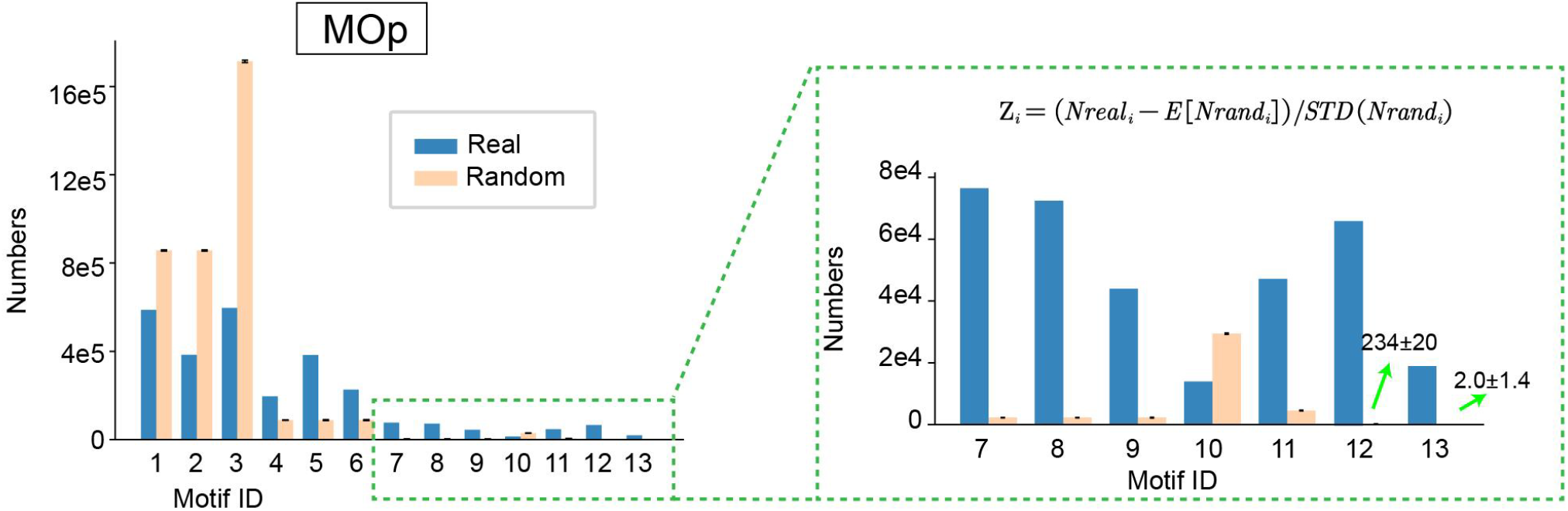
Number of three-node motifs in the MOp brain region versus matched ER random networks. Diagram showing counts of the 13 three-node motifs observed in the MOp region and the mean ± standard deviation of counts for each motif in the corresponding ER random network. These values were used to compute the normalized Z score for motif over- or under-representation.

**Fig. S8.**
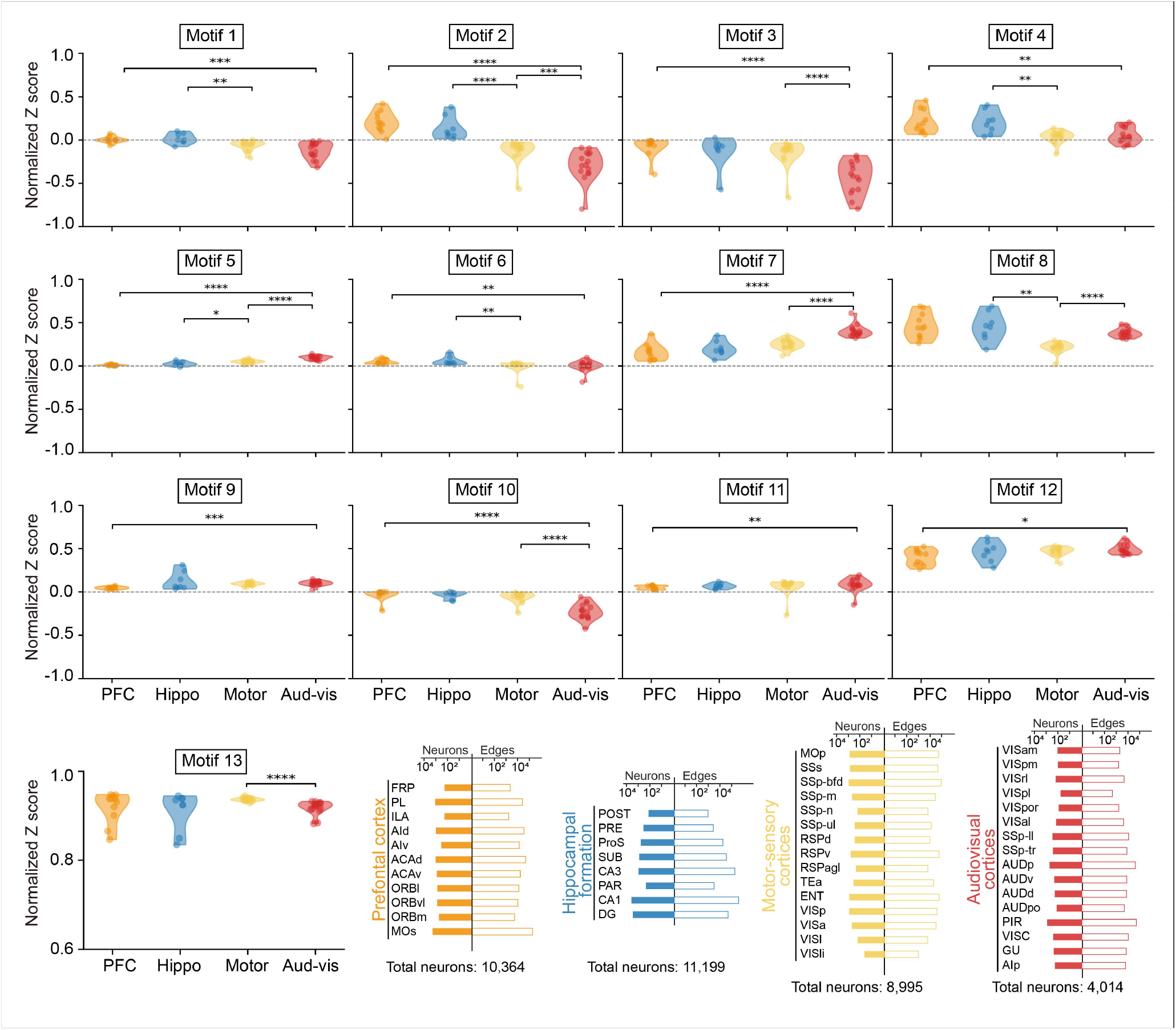
Frequency distributions of three-node motifs across brain region classes. Motif frequencies for all 13 three-node motifs across prefrontal cortex, hippocampal formation, motor-sensory cortices, and audiovisual cortices. Violin plots show normalized Z scores for each motif across the four brain region classes, with significance bars indicating pairwise differences among classes. The bottom panels summarize the included subregions together with their neuron and edge counts for each brain region class.

**Fig. S9.**
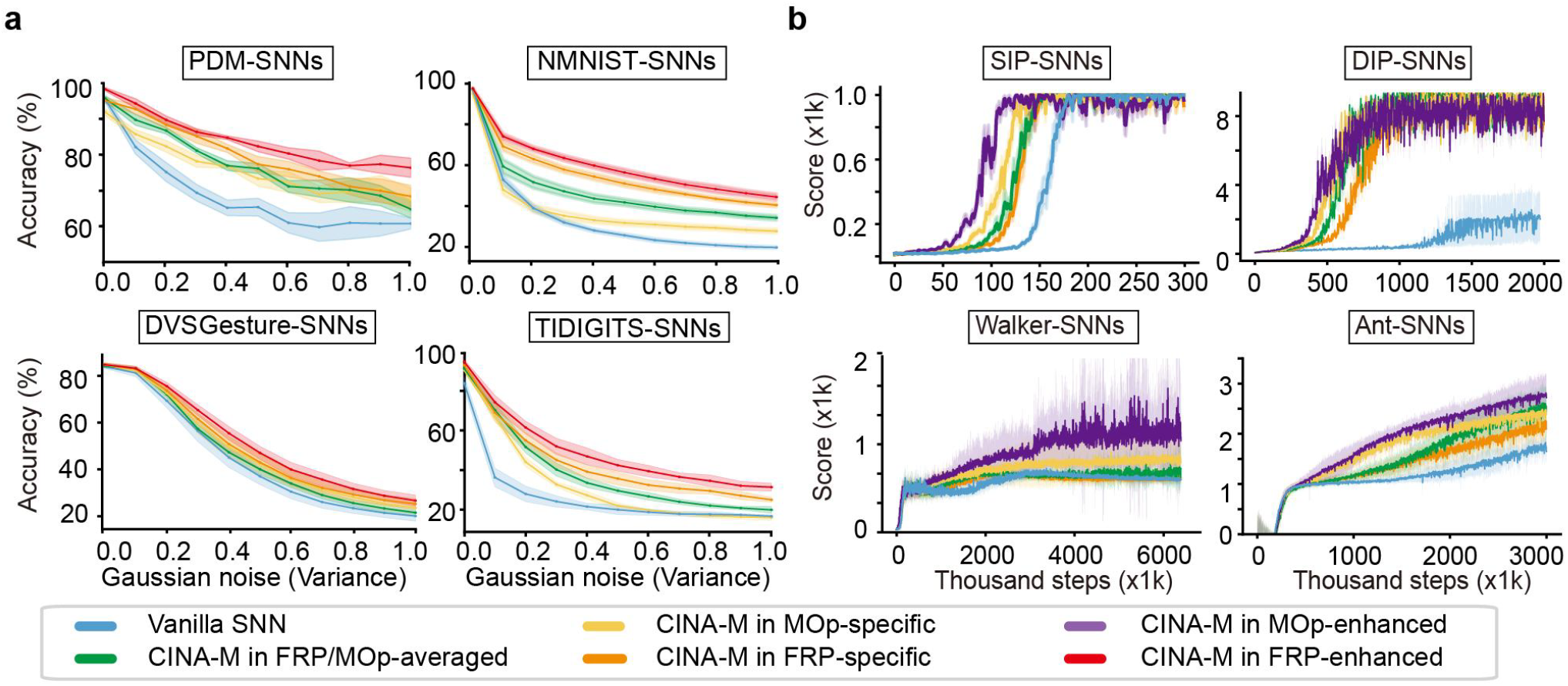
CINA-endowed biological and artificially biased motif distributions improve task performance of SNNs. **a**, Performance of CINA-endowed SNNs with the “vanilla” control in the four noise-resistant categorization tasks, shown as accuracy under increasing Gaussian noise variance. **b**, Performance of CINA-endowed SNNs with the “vanilla” control in the four motor learning tasks, shown as reward score during reinforcement learning. Different motif distributions are color-coded (box below).

**Fig. S10.**
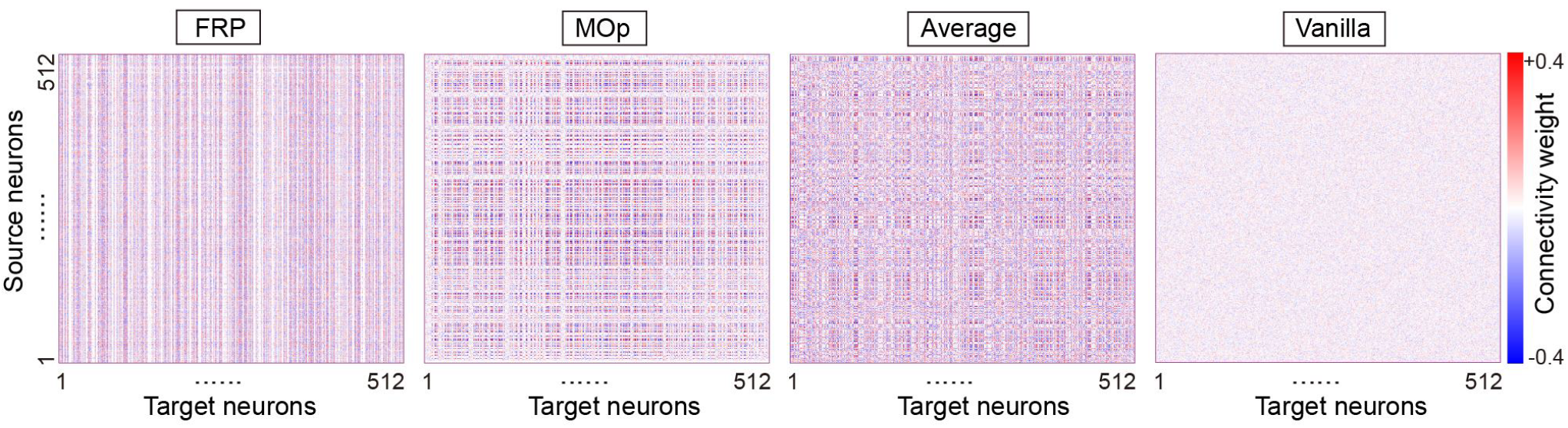
Connectivity weights of artificial neural networks. **a**, Heatmap diagram depicting connectivity weights of hidden pool neurons for ANNs endowed with “FRP-specific”, “MOp-specific”, “FRP/MOp-averaged” motif distributions and the “vanilla” distribution.

**Fig. S11.**
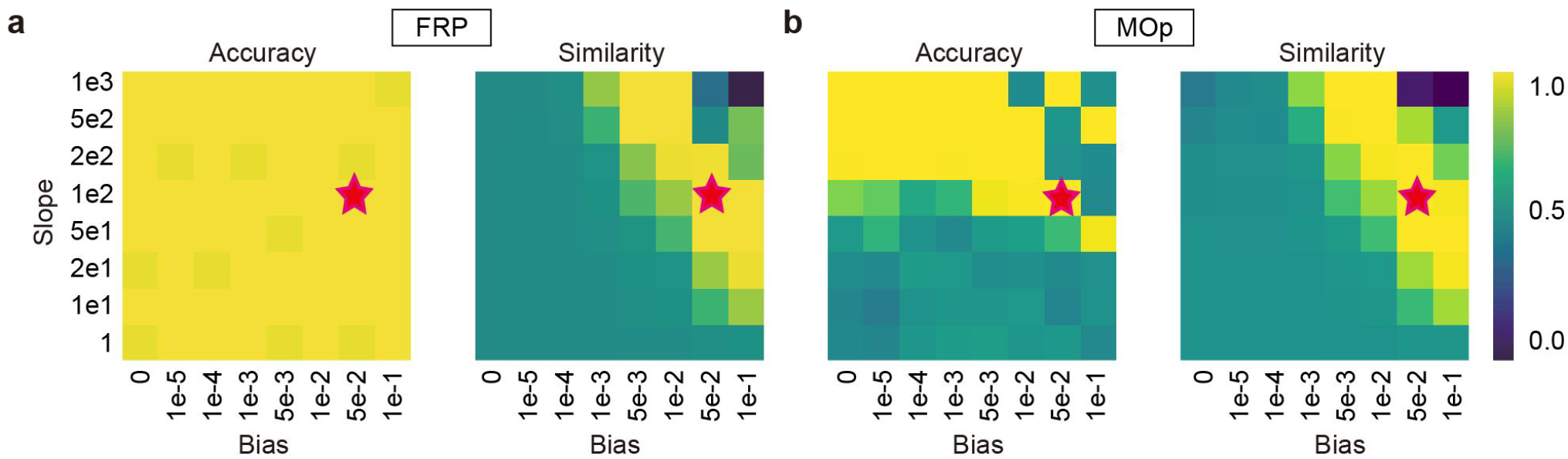
Accuracy and motif similarity of ANNs with FRP and MOp motif distributions. **a-b**, Accuracy and motif similarity of ANNs endowed with FRP (a) and MOp (b) motif distributions on a perceptual decision-making task across varying parameter configurations. The best configuration (Sigmoid bias = 0.05, slope = 100; red star) was selected for all tasks.

## SUPPLEMENTARY TABLES

**Table S1.**
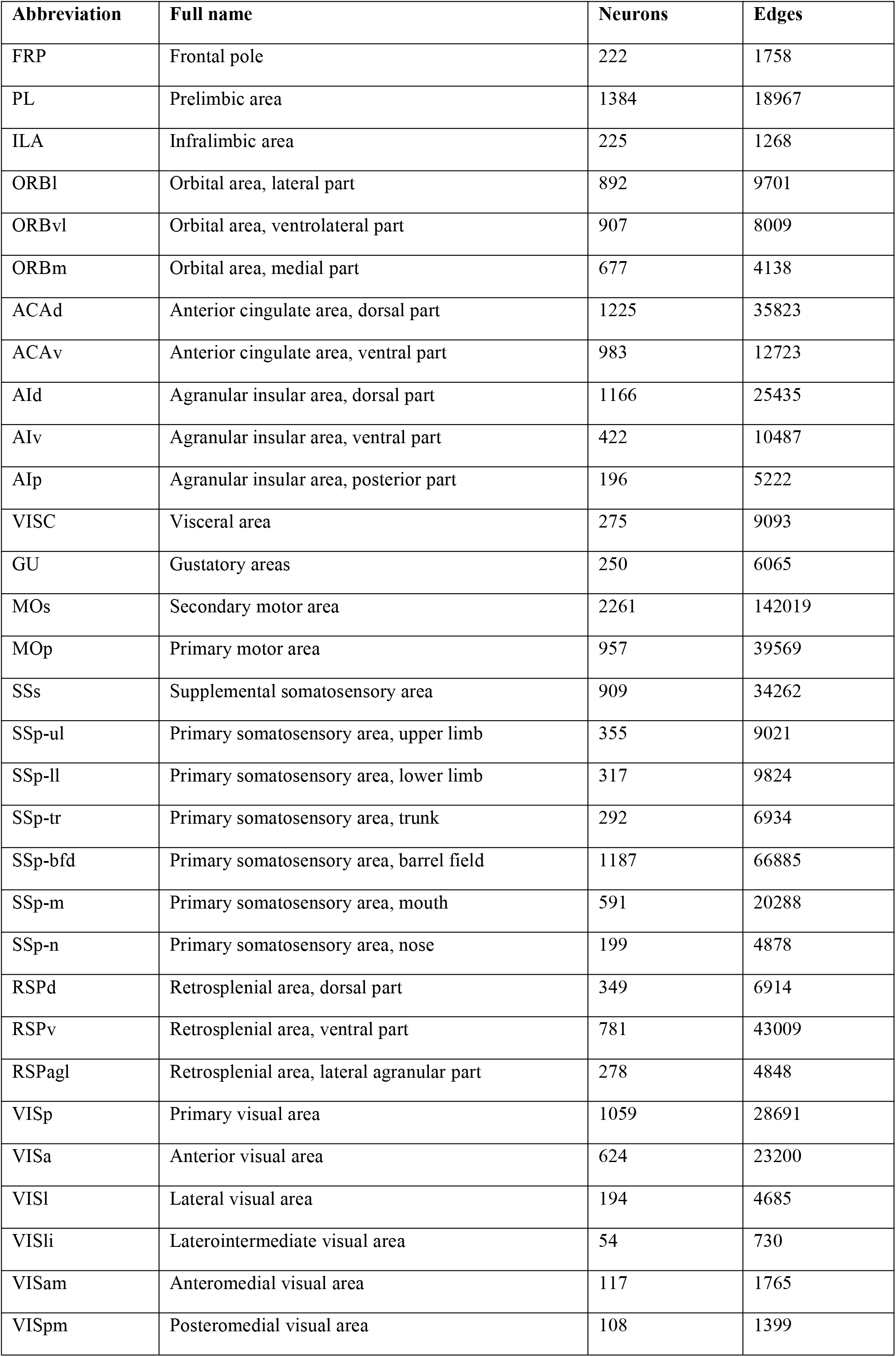
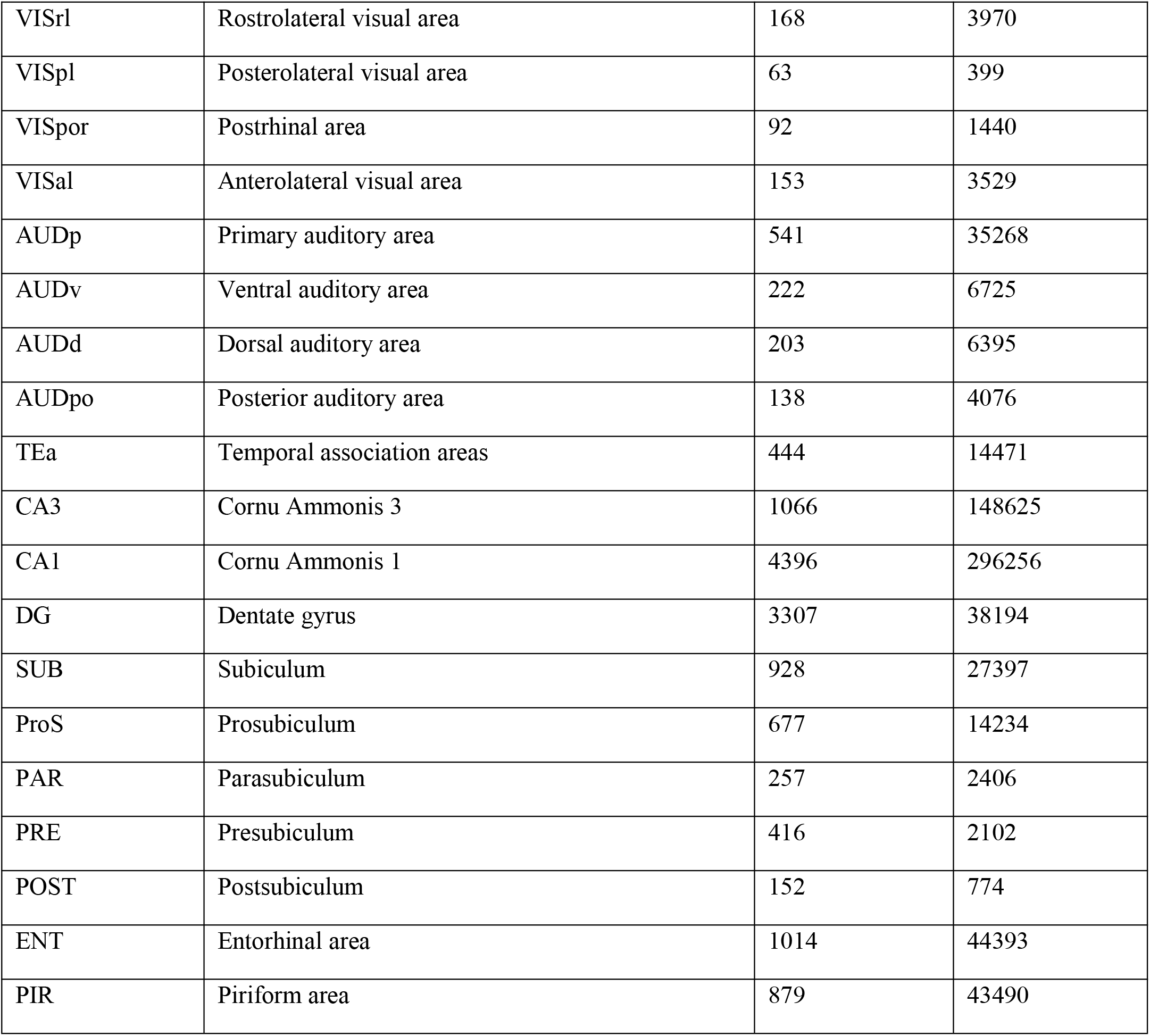
Number of neurons and synaptic connections in 50 brain regions.

**Table S2.**
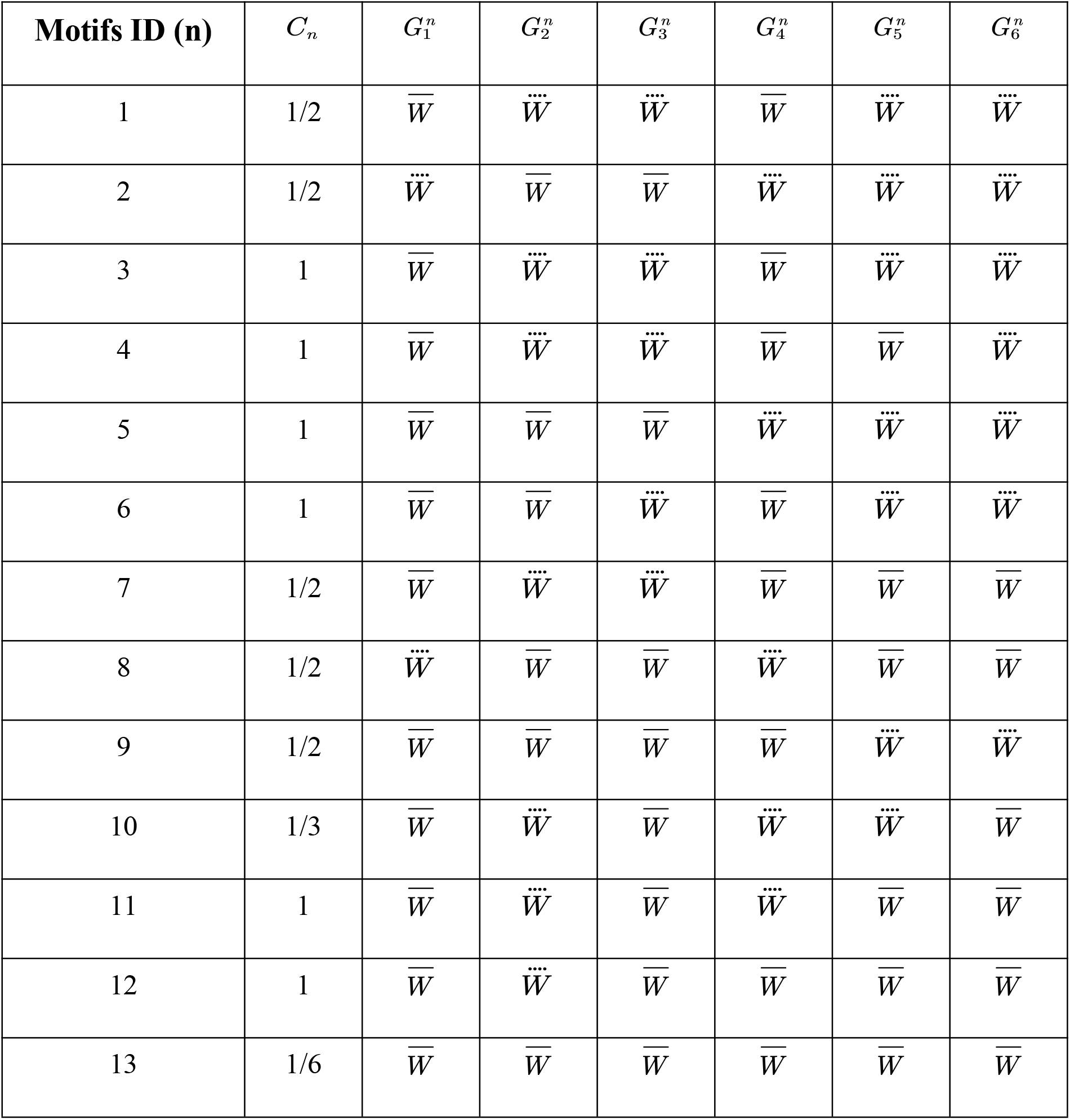
Variable values for 13 three-node motif formulas in Equation (2).

**Table S3.**
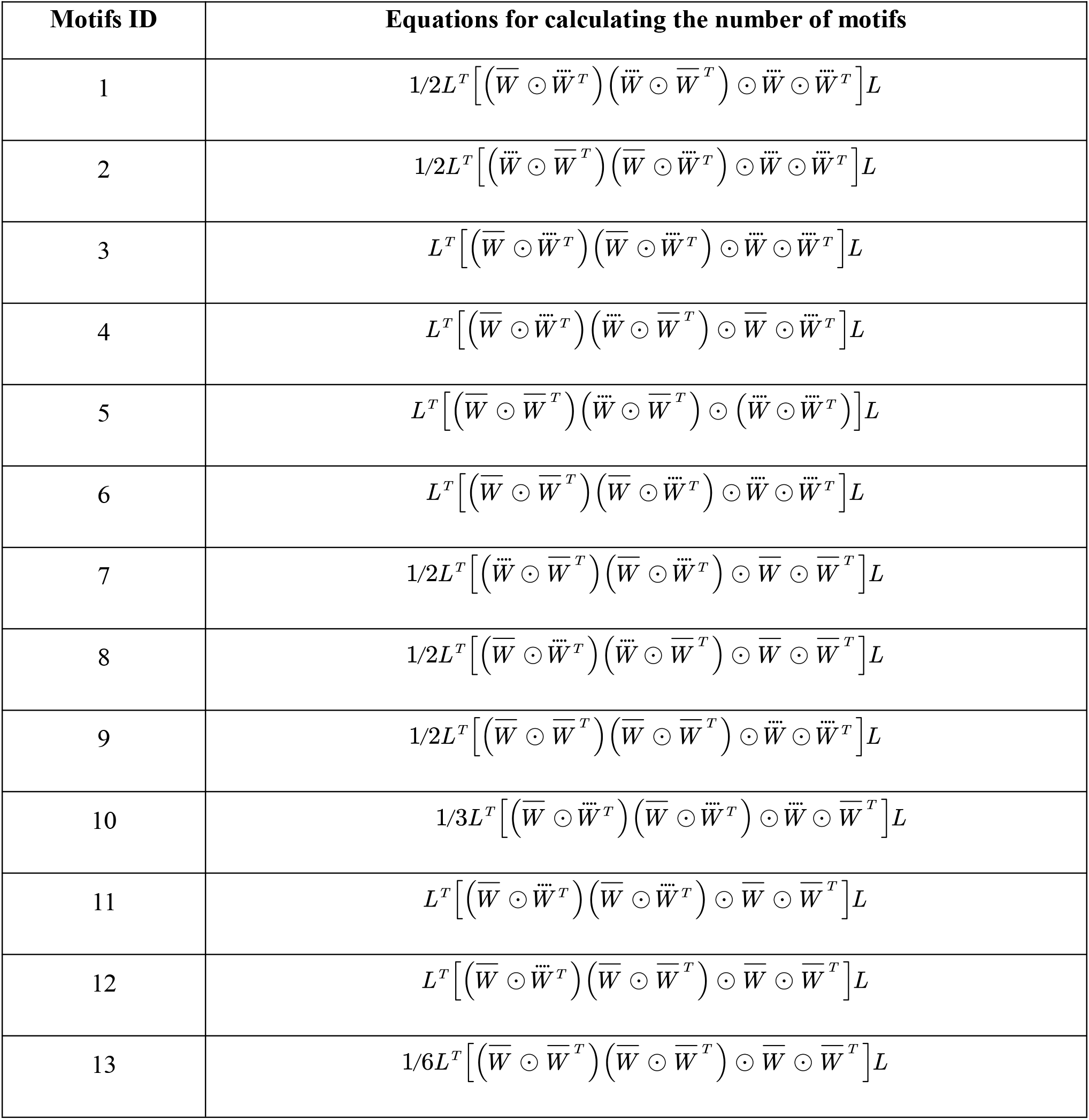
Equations for calculating the number of all 13 three-node motifs.

**Table S4.** Network configurations of different ANNs on benchmark recognition and RL tasks.

| Hyperparameter | PDM | MNIST | DVS<br>Gesture | TIDIGITS | Inverted<br>pendulum | Inverted<br>double<br>pendulum | Walker | Ant |
| --- | --- | --- | --- | --- | --- | --- | --- | --- |
| Input dimension | 1 | 28 | 256 | 20 | 4 | 9 | 17 | 27 |
| Hidden dimension | 512 | 512 | 512 | 256 | 512 | 512 | 512 | 512 |
| Output dimension | 1 | 10 | 11 | 10 | 1 | 1 | 6 | 8 |
| Batch size | 64 | 256 | 64 | 16 | 64 | 64 | 64 | 64 |
| Learning rate | 1e-3 | 1e-3 | 1e-3 | 1e-2 | 3e-5 | 3e-5 | 3e-5 | 3e-5 |
| Motif learning rate | 1e-2 | 1e-2 | 1e-2 | 1e-2 | 1e-2 | 1e-2 | 1e-2 | 1e-2 |
| Training epochs | 50 | 5 | 300 | 150 | 60 | 2000 | 4000 | 3000 |

